# Decoupling of spontaneous and evoked activity in locus coeruleus neurons: early synaptic and ion-channel remodeling in a mouse model of Alzheimer’s disease

**DOI:** 10.64898/2026.10.07.757306

**Authors:** Baizhen Zhao, Morgan Li, Lingxiao Yi, Jiali Li

## Abstract

The locus coeruleus (LC) is among the earliest brain regions affected in Alzheimer’s disease (AD), yet the mechanisms underlying its early vulnerability remain poorly understood. Clarifying synaptic and intrinsic dysfunction in LC noradrenergic (NE) neurons is essential to determine how noradrenergic circuit failure contributes to prodromal AD. We combined whole-cell patch-clamp recordings with quantitative immunofluorescence to examine synaptic transmission, firing properties, voltage-gated currents, and synaptic organization in LC/NE neurons from young 3xTg mice prior to substantial neurodegeneration. Young 3xTg mice exhibited reduced excitatory and enhanced inhibitory synaptic input onto LC/NE neurons. Consistent with these functional shifts, structural analysis revealed decreased densities of VGLUT2, PSD95, and paired VGLUT2–PSD95 puncta, alongside increased VGAT, gephyrin, and paired

VGAT – gephyrin puncta. Despite this net synaptic suppression, LC/NE neurons displayed elevated spontaneous firing rates but a markedly reduced capacity to generate spike output during sustained depolarization. Voltage-clamp recordings revealed a diminished total voltage-gated K⁺ current density alongside reduced voltage-gated Na⁺ current density, without significant shifts in their voltage-dependence or kinetics. These biophysical alterations provide a mechanistic basis for the divergent firing phenotype: reduced K⁺ conductance likely facilitates spontaneous pacemaker activity, whereas diminished Na⁺ current constrains repetitive, input-driven spike generation. Together, these findings demonstrate that LC/NE neurons undergo coordinated synaptic and ion-channel remodeling during prodromal AD. This unique decoupling of spontaneous baseline activity from stimulus-evoked excitability may impair dynamic noradrenergic modulation, positioning early LC cellular dysfunction as a candidate driver of network instability and a novel therapeutic target.

**Graphical Abstract:** 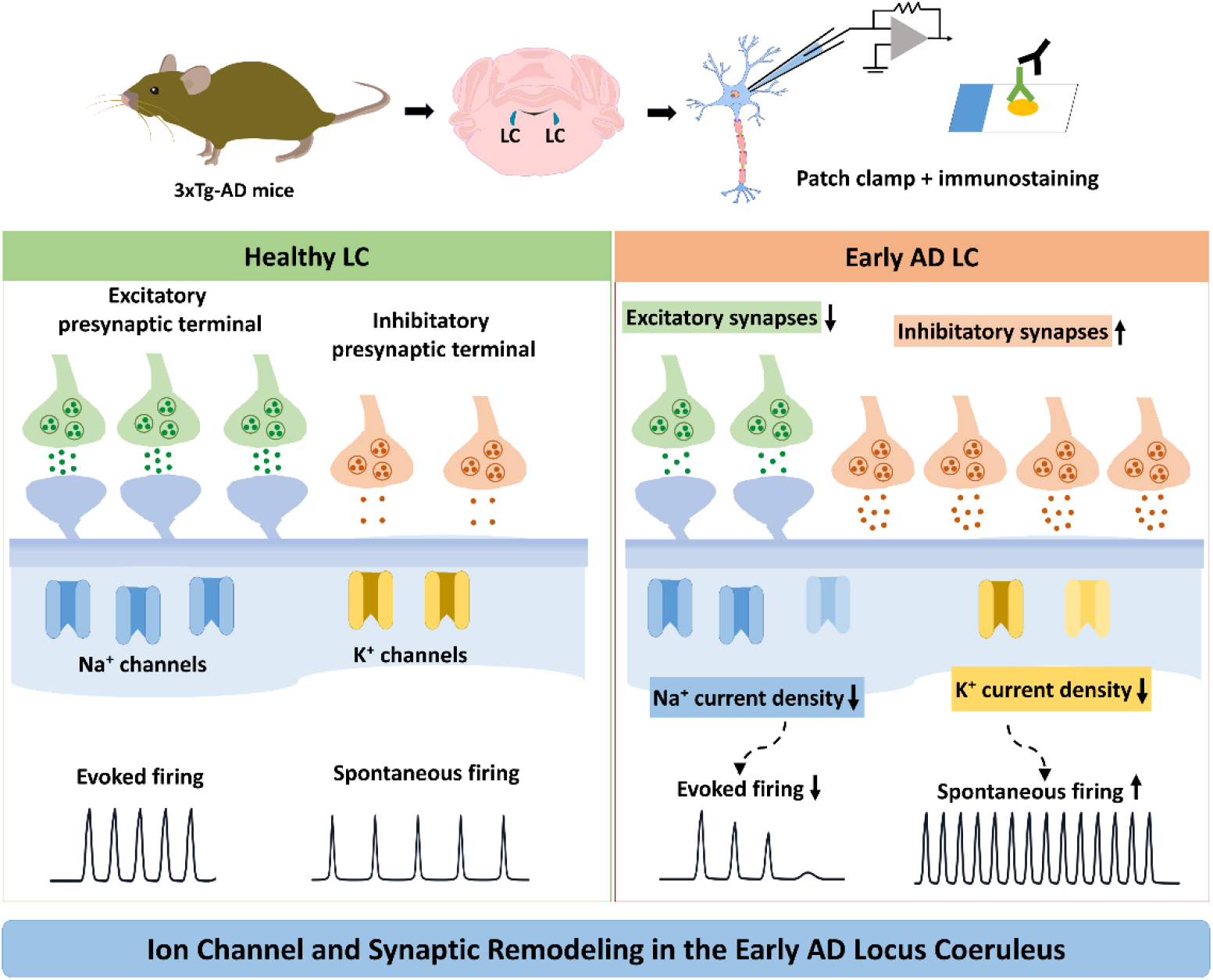

## Introduction

Alzheimer’s disease (AD) is characterized by progressive cognitive decline and the accumulation of amyloid-β (Aβ) plaques and neurofibrillary tau tangles, leading to widespread neuronal damage and brain atrophy(DeTure and Dickson, 2019; Goedert and Crowther, 1989; Goedert et al., 1991; Serrano-Pozo et al., 2011). Pathological changes begin years before overt cognitive symptoms, as shown by longitudinal imaging, fluid biomarker, and neuropathological studies (Bateman et al., 2012; Dubois et al., 2016; Forlenza et al., 2010; Jia et al., 2024; Rafii and Aisen, 2023; Thal et al., 2013; Villemagne et al., 2013). During this prodromal window, neuropsychiatric symptoms (NPS) such as anxiety, depression, apathy, irritability, and sleep disturbance frequently precede cognitive decline and predict faster progression to dementia (Finkel et al., 1996; Lyketsos et al., 2002; Nowrangi et al., 2015; Zhang et al., 2022), affecting over 90% of patients, yet their neural basis remains poorly understood.

The locus coeruleus (LC), a small brainstem nucleus of ∼1,600 noradrenergic neurons per hemisphere in humans, is the brain’s sole source of forebrain norepinephrine (NE)(Chen et al., 2022; Engels-Domínguez et al., 2024; Hary et al., 2025; Sun et al., 2020). Through projections to cortex, hippocampus, amygdala, and other limbic structures, the LC-NE system modulates arousal, attention, memory, emotion, and stress responses(Schwarz and Luo, 2015), functions commonly disrupted in AD(Bouret and Sara, 2005; Feinstein et al., 2016; Giorgi et al., 2020; Reyes, 2025). Notably, the LC accumulates hyperphosphorylated tau even in young, cognitively normal individuals(Braak and Del Tredici, 2011; Braak et al., 2011; Ehrenberg et al., 2017), , often decades before cortical tau pathology appears, suggesting it may be an initial site of AD-related change, with LC dysfunction potentially contributing to both early NPS and later cognitive decline through loss of noradrenergic modulation of downstream circuits. Despite this early vulnerability, the LC remains understudied relative to hippocampus and cortex, and the mechanisms linking LC dysfunction to AD pathogenesis are poorly defined. Neuromelanin-sensitive MRI studies show LC structural alterations in prodromal AD and mild cognitive impairment that correlate with tau burden and cognitive performance(Dahl et al., 2022), but the cellular and synaptic mechanisms underlying this vulnerability, and how they translate to circuit-level dysfunction and behavior, remain largely unknown.

Synaptic dysfunction is among the earliest events in AD pathogenesis, preceding substantial neuronal loss and cognitive decline(Scheff et al., 2007; Selkoe, 2002; Terry et al., 1991), and correlates more strongly with cognitive impairment than plaque or tangle burden(Colom-Cadena et al., 2020; Kazim et al., 2017; Kerchner and Nicoll, 2008; Koch et al., 2012; Koffie et al., 2011; Llinás, 2014; Mango et al., 2019; Matchett et al., 2021; Mercan and Heneka, 2022; Nadim and Bucher, 2014; Palop and Mucke, 2010; Prange and Murphy, 1999; Rafii and Aisen, 2023; Reyes, 2025; Robertson, 2013; Sara, 2009; Scheff et al., 2007; Schwarz and Luo, 2015; Selkoe, 2002; Serrano-Pozo et al., 2011; Spires-Jones and Hyman, 2014; Terry et al., 1991), positioning synaptic impairment as a primary driver of network failure rather than a downstream consequence of proteinopathy(Frere and Slutsky, 2018; Palop and Mucke, 2010). Functional and electrophysiological studies show altered synaptic plasticity, including impaired long-term potentiation (LTP) and aberrant long-term depression (LTD), in hippocampus and cortex early in disease progression(Di Lorenzo et al., 2019; Koch et al., 2012; Mango et al., 2019; Viana da Silva et al., 2016). Yet despite converging neuropathological evidence of even earlier pathology in the LC(Beardmore et al., 2021; Engels-Domínguez et al., 2024; Hary et al., 2025), systematic studies of LC synaptic function remain scarce. Given the LC’s widespread influence on forebrain circuits governing functions disrupted early in AD, whether and how LC synaptic transmission is altered could reveal critical mechanisms underlying prodromal disease progression.

These gaps lead to three key questions: (1) whether excitatory and inhibitory synaptic inputs onto LC neurons are altered early in AD, (2) whether such synaptic remodeling is accompanied by parallel changes in intrinsic firing properties and voltage-gated ionic conductance, and (3) how these synaptic and ionic alterations together shape LC output relevant to noradrenergic signaling and network stability in AD.

## Materials and Methods

### Animals

Transgenic mice of either sex aged 2–6 months were used as models of Alzheimer’s disease, including the 3xTg line (B6;129-Tg(APPSwe,tauP301L)1Lfa Psen1tm1Mpm/Mmjax) and age-matched wild-type littermate controls. All mouse lines were maintained on a C57BL/6 genetic background. Mice were group-housed in cohorts of three to five animals under specific pathogen–free conditions with ad libitum access to food and water. Animals were maintained on a 12 h light/dark cycle, with temperature (21–25°C) and humidity (45–55%) kept constant. All animal procedures were reviewed and approved by the Hackensack Meridian Health Animal Care and Use Committee at the Center for Discovery and Innovation (CDI) under protocol AUP-24-27 and conducted in accordance with institutional guidelines and applicable regulations.

### Acute brain slice preparation

Mice were deeply anesthetized with 3% isoflurane (induced in chamber, maintained via nose cone) and transcardially perfused with ice-cold, carbogen-saturated (95% O₂/5% CO₂) N-methyl-D-glucamine (NMDG)-based protective recovery solution containing (in mM): 92 NMDG, 2.5 KCl, 1.25 NaH₂PO₄, 30 NaHCO₃, 20 HEPES, 25 D-glucose, 2 thiourea, 5 sodium ascorbate, 3 sodium pyruvate, 0.5 CaCl₂·2H₂O, and 10 MgSO₄·7H₂O (osmolality 300-310 mOsm/kg, adjusted to pH 7.3-7.4 with HCl). Following decapitation, brains were rapidly extracted and immersed in ice-cold oxygenated NMDG solution. Coronal brainstem slices (300 μm thickness) containing the locus coeruleus (LC) were sectioned using a vibrating microtome (VT1200S, Leica Biosystems, Wetzlar, Germany) at a blade speed of 0.1 mm/s and vibration amplitude of 1.0 mm. Slices were immediately transferred to oxygenated NMDG solution at 34°C for 15 minutes to facilitate neuronal recovery, then transferred to carbogen-saturated artificial cerebrospinal fluid (ACSF) containing (in mM): 92 NaCl, 2.5 KCl, 1.25 NaH₂PO₄, 30 NaHCO₃, 20 HEPES, 25 D-glucose, 2 thiourea, 5 sodium ascorbate, 3 sodium pyruvate, 2 CaCl₂·2H₂O, and 2 MgSO₄·7H₂O (osmolality 300-310 mOsm/kg, adjusted to pH 7.3-7.4 with NaOH before oxygenation) and maintained at room temperature (22-24°C) for at least 60 minutes before recording. All solutions were continuously bubbled with carbogen (95% O₂/5% CO₂) throughout preparation and recovery periods.

### Whole-cell patch-clamp recordings Recording configuration

Individual slices were transferred to a submerged recording chamber mounted on an upright microscope (BX51WI, Olympus) and continuously superfused (2-3 mL/min) with carbogen-saturated recording ACSF containing (in mM): 124 NaCl, 2.5 KCl, 1.25 NaH₂PO₄, 24 NaHCO₃, 12.5 D-glucose, 5 HEPES, 2 CaCl₂·2H₂O, and 2 MgSO₄·7H₂O (osmolality 300-310 mOsm/kg, pH 7.3-7.4). LC neurons were visually identified using infrared differential interference contrast (IR-DIC) optics based on anatomical location lateral to the fourth ventricle at the level of the rostral pons and characteristic large soma. Recording pipettes were pulled from thick-walled borosilicate glass capillaries (1.5 mm outer diameter, 0.86 mm inner diameter; Sutter Instrument, BF150-86-10) using a horizontal puller (P-97, Sutter Instrument) and fire-polished to final resistances of 2-5 MΩ when filled with internal solution. Whole-cell recordings were obtained using a MultiClamp 700B amplifier, digitized with a Digidata 1550B interface, and acquired using pClamp 11 software (Molecular Devices, San Jose, CA). Signals were sampled at 10 kHz and low-pass filtered at 2 kHz using a 4-pole Bessel filter. Series resistance was monitored throughout recordings and cells were excluded from analysis if series resistance exceeded 25 MΩ or changed by >20% during recording.

### Voltage-clamp recordings

For spontaneous excitatory postsynaptic current (sEPSC) recordings, pipettes were filled with a potassium gluconate-based internal solution containing (in mM): 145 K-gluconate, 0.5 EGTA, 2 MgCl₂, 10 HEPES, 2 Mg-ATP, and 0.2 Na-GTP (osmolality 290-300 mOsm/kg, pH 7.3-7.4, adjusted with KOH). Neurons were voltage-clamped at-60 mV (uncorrected for liquid junction potential), and picrotoxin (100 μM, Tocris) was bath-applied to block GABA_A receptor-mediated currents. For miniature EPSC (mEPSC) recordings, tetrodotoxin (TTX, 1 μM, Biotium) was added to the bath to block action potential-dependent release, and the internal solution was cesium-based containing (in mM): 117 Cs-Gluconate, 5 TEA-Cl, 2.8 NaCl, 0.4 EGTA, 20 HEPES, 2 Mg-ATP, 0.2 Na-GTP (osmolality 290-300 mOsm/kg, pH 7.3-7.4, adjusted with CsOH). For sEPSC, the numbers of cells and animals included at each age were as follows: 2 months, WT: n=27 cells, N=6 mice and 3xTg: n=24 cells, N=5 mice; 4 months, WT: n=34 cells, N=8 mice and 3xTg: n=25 cells, N=5mice; and 6 months, WT: n=30 cells, N=6 mice and 3xTg: n=37 cells, N=7mice. For mEPSC, the numbers of animals included at each age were as follows: 2 months, WT: n=45 cells, N=10 mice and 3xTg: n=28 cells, N=7 mice; 4 months, WT: n=37 cells, N=7 mice and 3xTg: n=25 cells, N=5 mice; and 6 months, WT: n=18 cells, N=5 mice and 3xTg: n=18 cells, N=5 mice. Individual mice were considered biological replicates and individual cells were considered technical replicates.

For spontaneous inhibitory postsynaptic current (sIPSC) recordings, pipettes were filled with a high-chloride internal solution to enhance IPSC driving force, containing (in mM): 135 KCl, 5 EGTA, 5 MgCl₂, 10 HEPES, 2 Mg-ATP, and 0.2 Na-GTP (osmolality 290-300 mOsm/kg, pH 7.3-7.4, adjusted with KOH). Neurons were voltage-clamped at −60 mV in the presence of D-AP5 (50 μM, Tocris) and CNQX (10 μM, Tocris) to block NMDA and AMPA receptors, respectively. For miniature IPSC (mIPSC) recordings, TTX (1 μM) was bath-applied. sIPSCs and mIPSCs were recorded for 5-10 minutes per cell after establishing stable baseline. For sIPSC, the numbers of cells and animals included at each age were as follows: 2 months, WT: n=30 cells, N=5 mice and 3xTg: n=23 cells, N=5 mice; 4 months, WT: n=24 cells, N=4 mice and 3xTg: n=22 cells, N=4 mice; and 6 months, WT: n=24 cells, N=5 mice and 3xTg: 17 cells, N=5 mice. For mIPSC, the numbers of cells and animals included at each age were as follows: 2 months, WT: n=20 cells, N=5 mice and 3xTg: n=24 cells, N=5 mice; 4 months, WT: n=24 cells, N=5 mice and 3xTg: n=20 cells, N=5 mice; and 6 months, WT: n=12 cells, N=4 mice and 3xTg: n=20 cells, N=5 mice. Individual mice were considered biological replicates and individual cells were considered technical replicates.

For voltage-gated K^+^ current recording, Patch pipettes were filled with a K⁺-based internal solution containing (in mM): 125 K-gluconate, 5 KCl, 10 HEPES, 0.5 EGTA, 10 Na₂-phosphocreatine, 4 Mg-ATP, 0.3 Na₃-GTP (pH adjusted to 7.2–7.3 with KOH and osmolality of approximately 285–295 mOsm/kg). TTX (1 µM) was added to the bath solution to block voltage-gated Na⁺ currents and CdCl₂ (100 µM) to suppress voltage-gated Ca²⁺ currents and Ca²⁺-activated K⁺ currents. Additionally, D-AP5 (50μ M) and CNQX (10 μM) were included to minimize spontaneous synaptic currents.

Capacitive and leak currents were corrected using P/N=2 protocol. Liquid-junction potentials were not corrected. For voltage-gated K^+^ current recording, 2 month old 3xTg mice and age matched WT mice were used, the numbers of cells and animals included were as follows: WT: n=20 cells, N=5 mice and 3xTg: n=17 cells, N=5 mice. Individual mice were considered biological replicates and individual cells were considered technical replicates.

Voltage-gated Na⁺ currents were recorded using a Cs⁺-based intracellular solution containing (in mM): 70 Cs-gluconate, 30 Na-gluconate, 10 TEA-Cl, 5 4-aminopyridine, 10 EGTA, 1 CaCl₂, 10 HEPES, 4 Mg-ATP, 0.3 Na₃-GTP, and 10 Na₂-phosphocreatine (pH adjusted to 7.2–7.3 with CsOH and osmolality of approximately 285–295 mOsm/kg). CdCl₂ was added to the bath solution to suppress voltage-gated Ca²⁺ currents, D-AP5 and CNQX were included to minimize spontaneous synaptic currents, and intracellular Cs⁺, TEA, and 4-aminopyridine were used to minimize contamination from K⁺ currents. Linear leak and capacitive currents were subtracted using a P/N protocol with two scaled hyperpolarizing pulses (P/N = 2). Liquid-junction potentials were not corrected. For voltage-gated Na⁺ current recording, 2 month old 3xTg mice and age matched WT mice were used, the numbers of cells and animals included were as follows: recovery kinetics, WT: n=17 cells, N=5 mice and 3xTg: n=15 cells, N=5 mice; Na⁺ current density and activation kinetics, WT: n=16 cells, N=5 mice and 3xTg: n=13 cells, N=4 mice; Na⁺ inactivation kinetics, WT: n=15 cells, N=5 mice and 3xTg: n=14 cells, N=5 mice. Individual mice were considered biological replicates and individual cells were considered technical replicates.

### Current-clamp recordings

For spontaneous firing and evoked action potential recordings, the potassium gluconate-based internal solution (identical to that used for sEPSC recordings) was employed. Spontaneous firing was recorded in current-clamp mode for 4-5 minutes after achieving stable whole-cell configuration. Frequency-current (F-I) relationships were determined by injecting 1-second depolarizing current steps from 0 to 200 pA in 20 pA increments with 30-second inter-sweep intervals. Action potential threshold, amplitude, and half-width were analyzed from responses to the first suprathreshold current step. For spontaneous firing recording, the numbers of cells and animals included at each age were as follows: 2 months, WT: n=33 cells, N=6 mice and 3xTg: n=16 cells, N=5 mice; 4 months, WT: n=33 cells, N=7 mice and 3xTg: n=31 cells, N=6 mice; and 6 months, WT: n=19 cells, N=5 mice and 3xTg: n=20 cells, N=5 mice. For evoked action potential recordings, the numbers of animals included at each age were as follows: 2 months, WT: n=34 cells, N=7 mice and 3xTg: n=33 cells, N=7 mice; 4 months, WT: n=31 cells, N=7 mice and 3xTg: n=27 cells, N=7 mice; and 6 months, WT: n=25 cells, N=6 mice and 3xTg: n=30 cells, N=7 mice. Individual mice were considered biological replicates and individual cells were considered technical replicates.

### Data acquisition and inclusion criteria

All recordings were performed within 4-6 hours of slice preparation. Cells were included in analysis only if they met the following criteria: (1) stable resting membrane potential more negative than −45 mV (for current-clamp recordings); (2) stable holding current <200 pA at −60 mV (for voltage-clamp recordings); (3) series resistance <25 MΩ with <20% change during recording; (4) action potential amplitude >50 mV from threshold (for current-clamp recordings); and (5) stable baseline synaptic activity without rundown during recording period. These criteria resulted in inclusion of >85% of attempted recordings.

### Immunohistochemistry and image analysis

Mice were deeply anesthetized and transcardially perfused with phosphate-buffered saline (PBS), followed by 4% paraformaldehyde (PFA) in PBS. Brains were removed, post-fixed in 4% PFA at 4°C, and then dehydrated in 30% sucrose in PBS until they sank. Brain tissues were embedded in OCT compound and quickly frozen in −80℃ freezer for 10min, and then sectioned coronally at 15um thickness using a cryostat (Leica, USA). Sections containing the locus coeruleus (LC) were collected for immunofluorescence staining. Brain slices were washed three times in PBS with each time for 5min, and then permeabilized/blocked in blocking solution containing 5% bovine serum albumin (BSA) and 0.6% Triton X-100 in PBS for 1.5 hour at room temperature. Slices were then incubated overnight at 4°C with the following primary antibodies diluted in blocking solution containing 1% BSA and 0.3% Triton X-100: anti-TH (Invitrogen, OPA1-04050, 1:500), anti-VGLUT2 (Millipore Sigma, AB2251, 1:1000), anti-PSD95 (Invitrogen, MA1045, 1:500), anti-VGAT (Alomone Labs, AGT-005-GP, 1:500), anti-gephyrin (Synaptic Systems, 147011, 1:500). After primary antibody incubation, sections were washed three times in PBS and incubated with the fluorophore-conjugated secondary antibodies for 1 hour at room temperature in the dark, including goat anti-rabbit Alexa Fluor 488 and goat anti–guinea pig Alexa Fluor 594.

Sections were then washed three times again in PBS, counterstained with DAPI in antifade mounting medium.

Brain sections containing the locus coeruleus (LC) were identified based on anatomical landmarks and tyrosine hydroxylase (TH) immunoreactivity. Images were acquired from comparable rostrocaudal levels of the LC in wild-type (WT) and 3xTg mice. For each animal, three sections were examined, and two or three non-overlapping fields were collected from each section, ensuring that equivalent numbers of images were obtained from each experimental group. Image acquisition was performed at room temperature (∼22°C) using a Leica DMi8 confocal microscope system controlled via Leica Application Suite X (LAS X) software. High-resolution imaging of synaptic structures was executed using a 63× oil-immersion objective lens (Numerical Aperture, NA = 1.40) operating at a digital zoom of 2.5×. For the analysis of putative excitatory synapses, sections immunolabeled for TH, vesicular glutamate transporter 2 (VGLUT2), and postsynaptic density protein 95 (PSD95) were imaged. For the analysis of putative inhibitory synapses, sections immunolabeled for TH, vesicular GABA transporter (VGAT), and gephyrin were imaged.

Fluorophores were acquired sequentially using appropriate spectral detection windows and laser lines to eliminate spectral bleed-through. Hardware settings—including laser power, detector/smart gain, offset, pinhole size (set at approximately 1.0 Airy unit), scan speed, image resolution, and all other acquisition parameters—were kept strictly constant across experimental groups within each staining experiment. Images containing saturated pixels were excluded from analysis. Three-dimensional image stacks were acquired at a 1024 × 1024-pixel resolution with a z-step size of 0.30 µm, covering a total optical depth of 8 µm. Images were captured specifically from TH-positive somatic and proximal dendritic regions. The selection of imaging fields was based entirely on the TH channel without reference to the VGLUT2, PSD95, VGAT, or gephyrin channels to eliminate selection bias.

Image analysis was performed on the original, uncompressed z-stacks using Fiji (ImageJ) software equipped with the 3D Suite plugin. Background fluorescence was subtracted using identical settings across all images. VGLUT2-and PSD95-positive puncta were detected independently within or adjacent to a constructed TH-positive mask. A putative excitatory synaptic contact was operationally defined as a VGLUT2-positive presynaptic punctum paired with a PSD95-positive postsynaptic punctum within a maximum three-dimensional edge-to-edge distance of 0.4 µm. Similarly, VGAT-and gephyrin-positive puncta were detected independently, and a putative inhibitory synaptic contact was defined as a VGAT-positive punctum paired with a gephyrin-positive punctum within the same predetermined spatial criterion. Colocalization and structural proximity were evaluated throughout the complete volume of the z-stack by an experimenter blinded to the mouse genotypes. Immunofluorescence analyses included N=5 WT and N=5 3xTg mice. For each mouse, 3 anatomically matched LC sections and 3-5 non-overlapping imaging fields per section were analyzed. Individual mice were considered biological replicates, whereas sections and imaging fields were considered technical or observational replicates. Measurements from multiple fields and sections were averaged to generate one value per mouse for statistical analysis.

### Quantification and statistical analysis Electrophysiological data analysis

Electrophysiological recordings were digitized at 10 kHz and analyzed offline using Clampfit 11.3 (Molecular Devices, San Jose, CA) and MiniAnalysis software (Synaptosoft, Fort Lee, NJ). Only neurons meeting quality control criteria (stable series resistance <25 MΩ with <20% change during recording, stable baseline holding current, resting membrane potential more negative than −45 mV for current-clamp recordings) were included in analysis.

### Synaptic event detection

Spontaneous and miniature EPSCs and IPSCs were detected using automated template-matching algorithms in MiniAnalysis with a detection threshold set at 3× root-mean-square (RMS) baseline noise. All automatically detected events were visually inspected and false-positive events (artifacts, overlapping events) were manually excluded. Event frequency (Hz) was calculated as the total number of detected events divided by recording duration. Event amplitude (pA) was measured as the peak current from baseline. Kinetic parameters (10-90% rise time, decay time constant from single exponential fits) were measured from averaged events with >50 events per cell. For each cell, event parameters were averaged across the entire recording period (5-10 minutes), and each cell was treated as a single biological replicate (n).

### Spontaneous firing analysis

Spontaneous action potential frequency was calculated from 4-5-minute current-clamp recordings by dividing the total number of spikes by recording duration. Firing regularity was assessed by calculating the coefficient of variation (CV) of inter-spike intervals.

### Frequency-current (F-I) relationship analysis

Action potential frequency was counted for each 1-second current injection step (0-200 pA in 20 pA increments). F-I curves were constructed by plotting firing frequency against injected current. Input resistance was calculated from the voltage response to small hyperpolarizing current steps (−20 pA) using Ohm’s law.

## Statistical analysis

### Sample Sizes and Replicates

The numbers of biological replicates (*N*, representing individual mice) and technical replicates (*n*, representing individual cells or imaging fields) are explicitly detailed within each corresponding figure legend. Across all experimental cohorts, sample sizes ranged from N = 3 to 10 WT mice and N = 3 to 10 3xTg mice. Each cell was treated as an independent technical replicate nested within its respective biological replicate (*N*, animal). For quantitative immunofluorescence, biological replicates consisted of *N* = 5 WT mice and *N* = 5 3xTg mice. Technical replicates consisted of 2–3 non-overlapping imaging fields examined across 3 sections per animal, yielding a total structural analysis of 10–15 independent fields of view per mouse line. All statistical analyses were performed using GraphPad Prism 9.5 (GraphPad Software, San Diego, CA) or IBM SPSS Statistics 25.0. Data are presented as mean ± standard deviation (SD) unless otherwise noted. Sample sizes (n = number of cells, N = number of animals) are reported in figure legends and represent biologically independent samples.

### Normality and variance testing

Data distributions were assessed for normality using the Shapiro-Wilk test. Homogeneity of variance was tested using Levene’s test for independent samples or Mauchly’s test of sphericity for repeated measures designs.

### Comparison tests

For comparisons between two groups (WT vs. 3xTg at single timepoints), unpaired two-tailed Student’s t-tests were used for normally distributed data; Mann-Whitney U tests were used for non-normally distributed data. For F-I curve analysis involving repeated current steps, two-way repeated measures ANOVA was performed with genotype as between-subjects factor and current injection as within-subjects factor, followed by Bonferroni post-hoc tests for multiple comparisons when main effects or interactions were significant (p < 0.05). Greenhouse-Geisser corrections were applied when sphericity assumptions were violated.

### Effect sizes

Cohen’s d was calculated for t-test comparisons and partial eta-squared (η²_p) for ANOVA to quantify effect sizes. Cohen’s d values of 0.2, 0.5, and 0.8 represent small, medium, and large effects, respectively.

### Statistical significance

Statistical significance was defined as p < 0.05. Exact p-values are reported in figures and text when p ≥ 0.001; for p < 0.001, this is indicated. Significance levels are denoted as: p < 0.05; *p < 0.01; **p < 0.001. All tests were two-tailed unless otherwise specified.

### Data visualization

Individual data points were overlaid on bar graphs to display the distribution of measurements across cells. Summary data are presented as mean ± SEM unless otherwise indicated. Each data point represents a single recorded neuron, with the number of cells (n) and animals (N) specified in the figure legends. Bar graphs were used to illustrate group means, while superimposed scatter plots provide transparency regarding data variability and sample size.

### Sample-size determination, statistical assumptions

Sample sizes were based on preliminary experiments and published studies using similar approaches in AD models. Individual mice were considered biological replicates (N), whereas cells, tissue sections, and non-overlapping imaging fields obtained from the same mouse were considered observational or technical replicates (n). Statistical inference was based on mouse-level values or analyses that accounted for observations nested within individual mice. Sample sizes are reported as n = cells/N = mice for electrophysiological experiments and as the numbers of mice, sections, and imaging fields for imaging experiments. Data distributions were assessed using the Shapiro–Wilk test, and homogeneity of variance was assessed using Levene’s test. Data satisfying both assumptions were analyzed using the indicated parametric tests. When the data were not normally distributed or the variances were heterogeneous, nonparametric tests were used. Comparisons between two independent groups were performed using the Mann–Whitney U test.

### Randomization and blinding

Animals were not randomized by genotype because genotype was determined by breeding. To minimize potential order and batch effects, WT and 3xTg animals were interspersed across experimental days, and samples from the two genotypes were processed in parallel or in alternating order whenever possible. Electrophysiological recordings and imaging were performed using predefined acquisition parameters and prespecified inclusion and exclusion criteria. For immunofluorescence quantification and statistical analysis, data files were coded so that investigators were blinded to genotype until completion of the analysis. Identical acquisition and analysis settings were applied across experimental groups. These procedures were designed to minimize subjective bias and improve the transparency and reproducibility of data collection and analysis, consistent with published recommendations for rigorous experimental design, statistical analysis, and data presentation(Michel et al., 2020; Weissgerber et al., 2015).

### Experimental design

Animals were assigned to experimental groups based on genetically determined genotype (3xTg-AD or WT littermates) and age (2, 4, or 6 months), with both sexes included and distributed approximately evenly across groups. Because genotype was determined by breeding, animals could not be randomized by genotype. To minimize potential order and batch effects, animals from the two genotypes were interspersed across experimental days, and samples were processed in parallel or in alternating order whenever possible. Experimenters performing electrophysiological recordings and initial data acquisition were blinded to genotype. Predefined acquisition parameters and inclusion/exclusion criteria were applied throughout the experiments. For immunofluorescence quantification and statistical analysis, data files were coded so that investigators remained blinded to genotype until completion of the analysis. Identical acquisition and analysis settings were applied across experimental groups.

## Results

### Hyperexcitability of LC/NE neurons in young 3xTg mice

Neuronal hyperexcitability represents one of the earliest pathophysiological hallmarks of AD, preceding the emergence of classical neuropathological markers and cognitive decline (Kazim et al., 2017; Yook et al., 2024). To investigate potential alterations in the excitability of LC neurons during the early stages of AD, we performed electrophysiological recordings of spontaneous firing activity in 3xTg mice at 2-, 4-, 6-month-old representing distinct phases of disease progression (Figure 1A, B), these ages represent the early stages of disease progression in the 3xTg model, occurring prior to the emergence of extensive amyloid plaques and neurofibrillary tangles but coinciding with early biochemical and functional alterations. Brainstem slice containing the LC was prepared with an NMDG-based slicing protocol. LC was identified by their position in relation to the fourth ventricle and characterized by dense packed large neurons. To ensure noradrenergic neurons (LC/NE neurons) were exclusively studied, we targeted the core region of LC, which is almost entirely composed of noradrenergic neurons(Caestecker et al., 2025; Chandler, 2016). The spontaneous firing rate serves as a fundamental metric of baseline neuronal excitability, integrating contributions from intrinsic membrane properties, synaptic inputs, and neuromodulatory influences(Alejandre-García et al., 2022; Jacob and Nienborg, 2018; Llinás, 2014; Nadim and Bucher, 2014; Yamada-Hanff and Bean, 2013).

**Figure 1.**
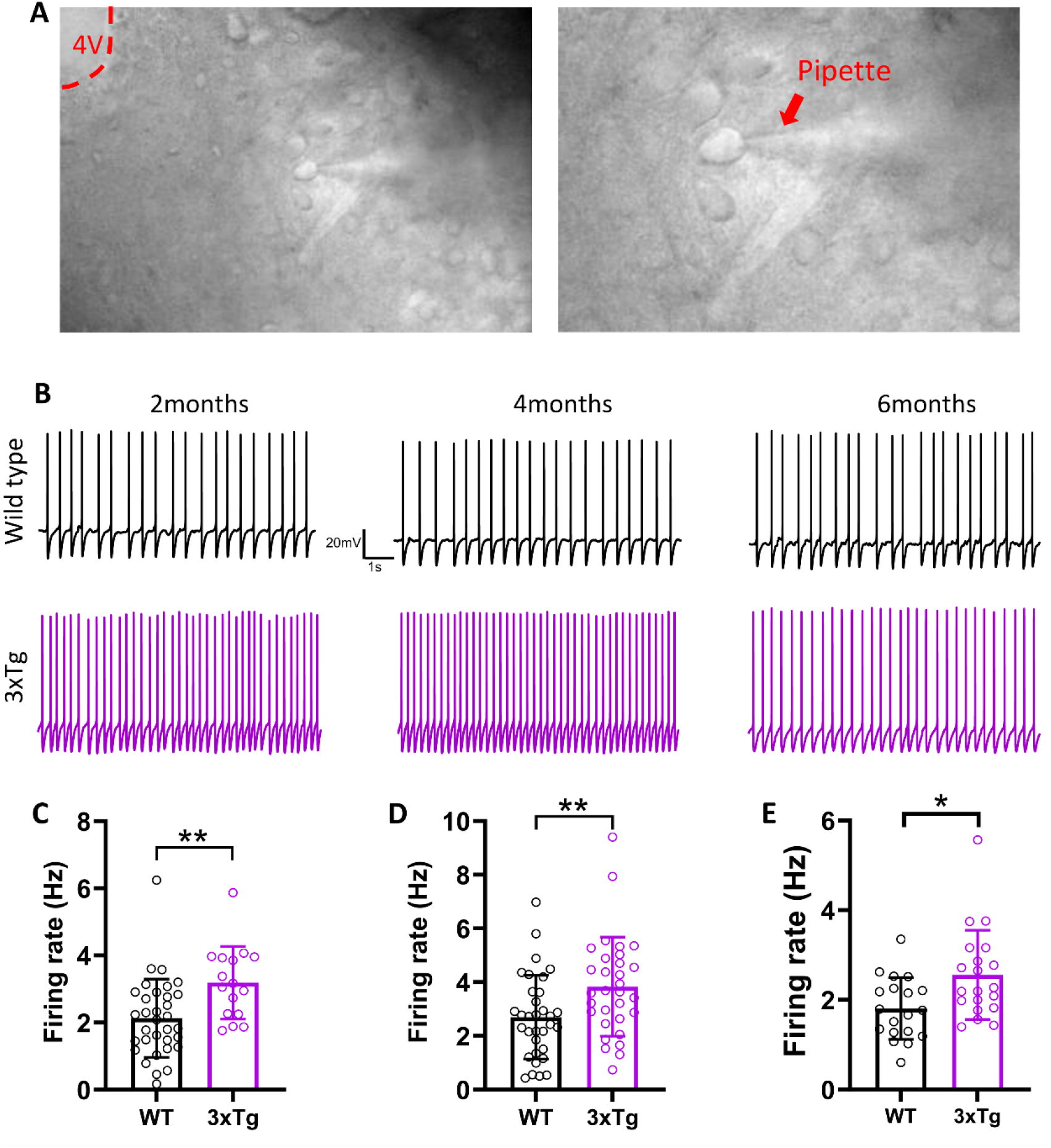
Hyperexcitability of LC/NE neurons in young 3xTg mice. **A** Coronal slices prepared from 3xTg mice for whole-cell patch-clamp recordings of LC/NE neurons. Left: location of the LC near the fourth ventricle (4v). Right: recording pipette approaching an LC neuron. **B** Representative traces of spontaneous firing recorded from LC neurons at different ages. **C** Spontaneous firing frequency of LC neurons at 2 months of age (WT: 2.13 ± 1.17 Hz, n = 33 cells, N = 6 mice; 3xTg: 3.19 ± 1.08 Hz, n = 16 cells, N = 5 mice; Shapiro – Wilk tests, WT: W=0.923, p=0.023, 3xTg: W=0.922, p=0.181; Levene’s test, F_(1,9)_=0.441, p=0.523; unpaired two-tailed Mann Whitney U test, U=117, p = 0.001). **D** Spontaneous firing frequency of LC neurons at 4 months of age (WT: 2.69 ± 1.56 Hz, n = 33 cells, N = 7 mice; 3xTg: 3.83 ± 1.84 Hz, n = 31 cells, N = 6 mice; Shapiro–Wilk tests, WT: W=0.953, p=0.17, 3xTg: W=0.938, p=0.073; Levene’s test, F_(1,11)_=0.122, p=0.733; unpaired two-tailed t test, t₆₂ = 2.672, p = 0.009). **E** Spontaneous firing frequency of LC neurons at 6 months of age (WT: 1.81 ± 0.69 Hz, n = 19 cells, N = 5 mice; 3xTg: 2.55 ± 0.99 Hz, n = 20 cells, N = 5 mice; Shapiro–Wilk tests, WT: W=0.972, p=0.81, 3xTg: W=0.876, p=0.015; Levene’s test, F_(1,8)_=2.341, p=0.165; unpaired two-tailed Mann Whitney U test, U=99, p = 0.009). Data are represented as mean ± SD. Each point represents an individual recorded cell. n, number of cells; N, number of animals. *p < 0.05; **p < 0.01.

At 2 months of age, LC neurons from 3xTg mice exhibited a markedly elevated spontaneous firing rate compared to age-matched wild-type (Figure 1C). This elevation in spontaneous firing represents a significant increase relative to WT controls, indicating substantial hyperexcitability of LC neurons at this early point. The hyperexcitable phenotype persisted at 4 months of age, with 3xTg mice continuing to display significantly increased spontaneous firing rates compared to WT controls (Figure 1D). The sustained elevation of firing rates at this intermediate age suggests that the hyperexcitable state established early in disease progression remains stable through the first several months of development in the 3xTg model. Notably, by 6 months of age, LC neurons from 3xTg mice continued to exhibit an elevated spontaneous firing rate (Figure 1E), indicating persistent hyperexcitability. However, the magnitude of this increase was reduced compared to that observed at 2 and 4 months, suggesting a partial attenuation of hyperexcitability at later stages. This change may reflect compensatory adaptations or a transition to a distinct pathophysiological state as the disease progresses.

### Reduction in evoked excitability of LC/NE neurons found in young 3xTg mice

While spontaneous firing reflects the integration of intrinsic properties with ongoing synaptic inputs and neuromodulatory tone, evoked firing patterns in response to defined current injections provide a direct assessment of intrinsic membrane excitability, largely independent of network activity. To this end, we performed systematic current-clamp recordings in which LC/NE neurons were subjected to incrementally increasing depolarizing current injections (0–200 pA in 20 pA steps), and we quantified the relationship between injected current amplitude and action potential firing frequency (frequency-current or F-I relationships) across multiple ages in both 3xTg and WT mice (Figure 2A). Contrary to what might be predicted based on the elevated spontaneous firing rates observed in young 3xTg mice, our evoked excitability experiments revealed a striking and unexpected phenotype. At 2 months of age, current-clamp recordings demonstrated that LC neurons from 3xTg mice generated significantly fewer action potentials in response to depolarizing current steps compared to WT neurons (Figure 2B). Representative traces illustrate the marked reduction in spike number at higher current intensities in 3xTg neurons relative to WT controls. Quantitative analysis of the complete F-I relationship confirmed that 3xTg neurons exhibited significantly suppressed evoked firing across the stimulus range. Notably, the divergence between genotypes became statistically significant at the 100 pA current step and persisted at all higher current intensities. This observation indicates that while LC neurons from 2-month-old 3xTg mice can generate action potentials in response to weak-to-moderate depolarizing inputs, their capacity to sustain high-frequency firing in response to strong depolarization is markedly impaired.

**Figure 2.**
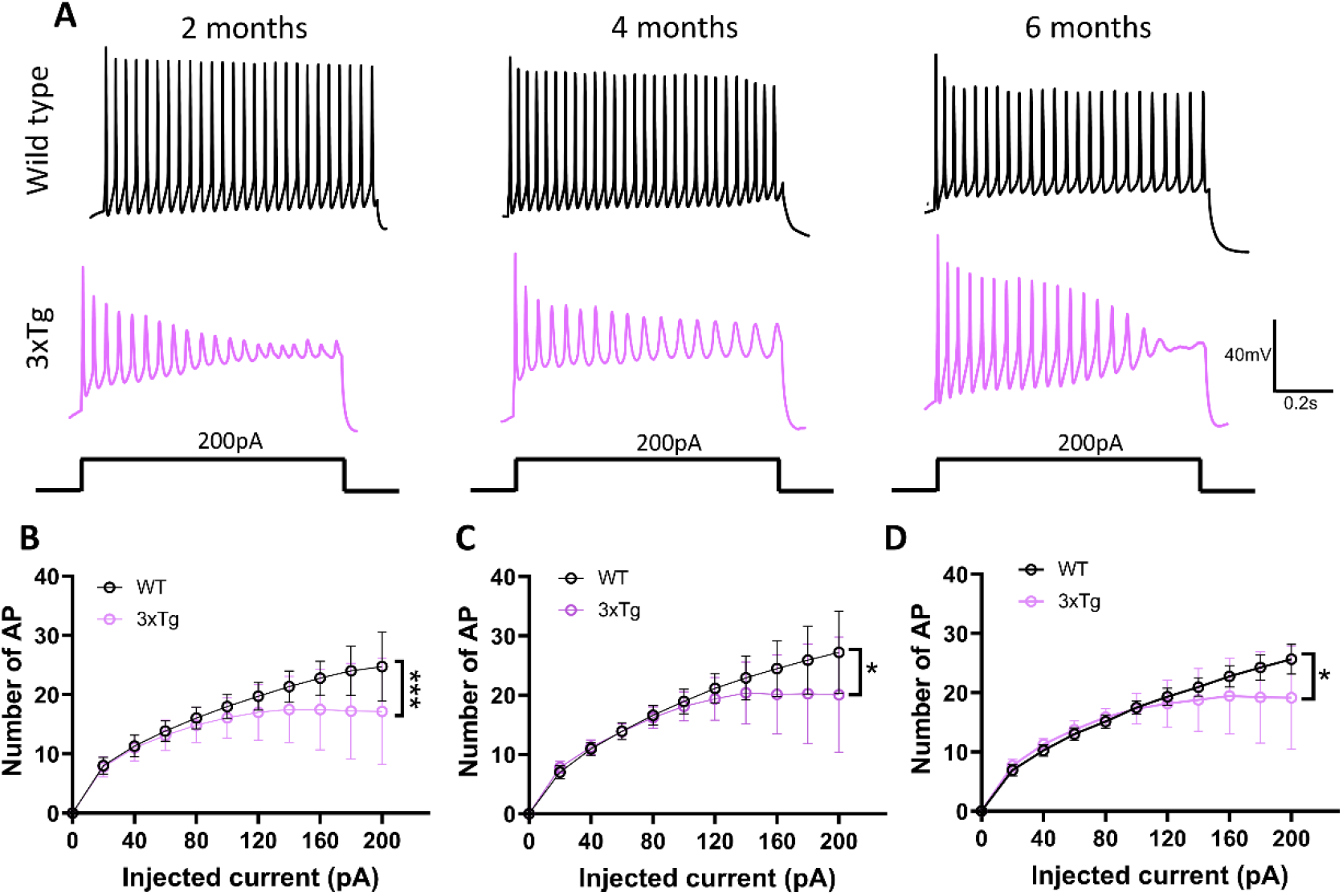
Reduction in evoked excitability of LC/NE neurons in young 3xTg mice. **A** Representative traces of action potentials evoked by current injection (0–200 pA, 20 pA increments) at different ages. **B** Action potential frequency of LC neurons at 2 months of age as a function of injected current (WT: n = 34 cells, N = 7 mice; 3xTg: n = 33 cells, N = 7 mice; two-way repeated-measures ANOVA, F₁, ₆₅ = 15.68, p < 0.001). **C** Action potential frequency of LC neurons at 4 months of age as a function of injected current (WT: n = 31 cells, N = 7 mice; 3xTg: n = 27 cells, N = 7 mice; two-way repeated-measures ANOVA, F₁, ₅₆ = 7.01, p = 0.010). (**D)** Action potential frequency of LC neurons at 6 months of age as a function of injected current (WT: n = 25 cells, N = 6 mice; 3xTg: n = 30 cells, N = 7 mice; two-way repeated-measures ANOVA, F₁, ₅₃ = 4.29, p = 0.040). Data are represented as mean ± SD. Individual values are not displayed at every current step to preserve the clarity of the input–output curves. n, number of cells; N, number of animals. *p < 0.05; **p < 0.01; ***p < 0.001.

The reduction in evoked excitability persisted at 4 months of age, with 3xTg neurons continuing to display significantly attenuated F-I relationships compared to WT controls (Figure 2C). Interestingly, at this age, the threshold at which genotype differences emerged shifted to the 120 pA current step, representing a slightly higher current intensity compared to the 100 pA threshold observed at 2 months. This rightward shift in the divergence point may suggest subtle age-dependent changes in the specific mechanisms underlying the reduced excitability phenotype, potentially reflecting progressive alterations in ion channel expression, distribution, or biophysical properties as disease pathology advances. Despite this shift in threshold, the overall magnitude of the deficit in evoked firing remained substantial, with 3xTg neurons generating significantly fewer spikes across the higher current range (120–200 pA). By 6 months of age, the reduced evoked excitability phenotype remained evident, although the statistical significance of the genotype effect was somewhat diminished compared to earlier ages (Figure 2D). The divergence between genotypes did not become apparent until the 160 pA current step, representing a further rightward shift compared to younger ages (100 pA at 2 months, 120 pA at 4 months). This progressive increase in the current threshold required to reveal genotype differences suggests that the intrinsic mechanisms underlying reduced excitability may be evolving with age, potentially becoming more selective for high-frequency firing regimes. The continued presence of reduced evoked excitability at 6 months (Figure 2D), indicates that these two aspects of neuronal excitability (spontaneous vs. evoked) are dissociable and likely governed by distinct cellular mechanisms or regulatory processes.

The temporal progression of the reduced evoked excitability phenotype reveals several important features of LC dysfunction in the 3xTg model. First, the deficits are present as early as 2 months of age and persist through at least 6 months, indicating that impaired intrinsic responsiveness is an early and sustained pathophysiological feature. Second, the progressive rightward shift in the current threshold at which genotype differences emerge (from 100 pA at 2 months to 160 pA at 6 months) suggests age-dependent refinement of the deficit, with older neurons showing vulnerability specifically during high-frequency firing demands. Third, the gradual reduction in the statistical significance of the genotype effect with age may reflect either increasing phenotypic variability within the 3xTg population, subtle compensatory mechanisms that partially ameliorate the deficit, or age-related changes in WT neurons that reduce the magnitude of genotype differences. Indeed, the coexistence of elevated spontaneous firing (Figure 1) with reduced evoked excitability (Figure 2) in LC/NE neurons from young 3xTg mice provides important mechanistic insights. This bidirectional alteration in excitability properties indicates that spontaneous and evoked firing are mediated by distinct biophysical mechanisms that are differentially affected in the 3xTg model. Spontaneous firing in LC neurons is primarily driven by persistent sodium currents, hyperpolarization-activated cation currents (Ih), and calcium-activated potassium currents, which together establish a depolarized resting membrane potential and enable autonomous pacemaker activity. In contrast, high-frequency evoked firing requires robust activation and rapid recovery of transient voltage-gated sodium channels, efficient repolarization mediated by delayed rectifier potassium channels, and effective regulation of intracellular calcium to prevent spike frequency adaptation.

### Impairment of glutamatergic transmission in LC/NE neurons of young 3xTg mice

To determine whether early synaptic dysfunction in LC/NE neurons occurs prior to overt neurodegeneration, we examined excitatory synaptic transmission, spontaneous and miniature postsynaptic currents (sEPSCs/mEPSCs) of LC/NE neurons at three critical developmental time points 2, 4, and 6 months of age were recorded. The combined analysis of sEPSCs and mEPSCs provides complementary information about distinct aspects of synaptic function. sEPSCs represent total excitatory input to a neuron at rest including both action potential-dependent and -independent neurotransmitter release, while mEPSCs specifically isolate action potential-independent vesicle fusion events. Changes in sEPSC frequency and amplitude thus primarily reflect alterations in presynaptic circuit activity and postsynaptic receptor strength, respectively, whereas changes in mEPSC frequency indicate modifications in vesicle release probability or synapse number, and changes in mEPSC amplitude reveal alterations in postsynaptic receptor number or conductance(Kavalali et al., 2011; Kerchner and Nicoll, 2008; Prange and Murphy, 1999; Turrigiano, 2008). sEPSCs was recorded at −60mV in the presence of GABA_A_ receptor inhibitor picrotoxin to isolate glutamatergic currents and block inhibitory transmission. Analysis of sEPSC frequency revealed a robust and persistent reduction in excitatory synaptic activity onto LC neurons in 3xTg mice across all three ages examined, indicating chronic suppression of glutamatergic circuit activity throughout the early disease period (Figure 3A). At 2 months of age, LC neurons from 3xTg mice exhibited significantly reduced

**Figure 3.**
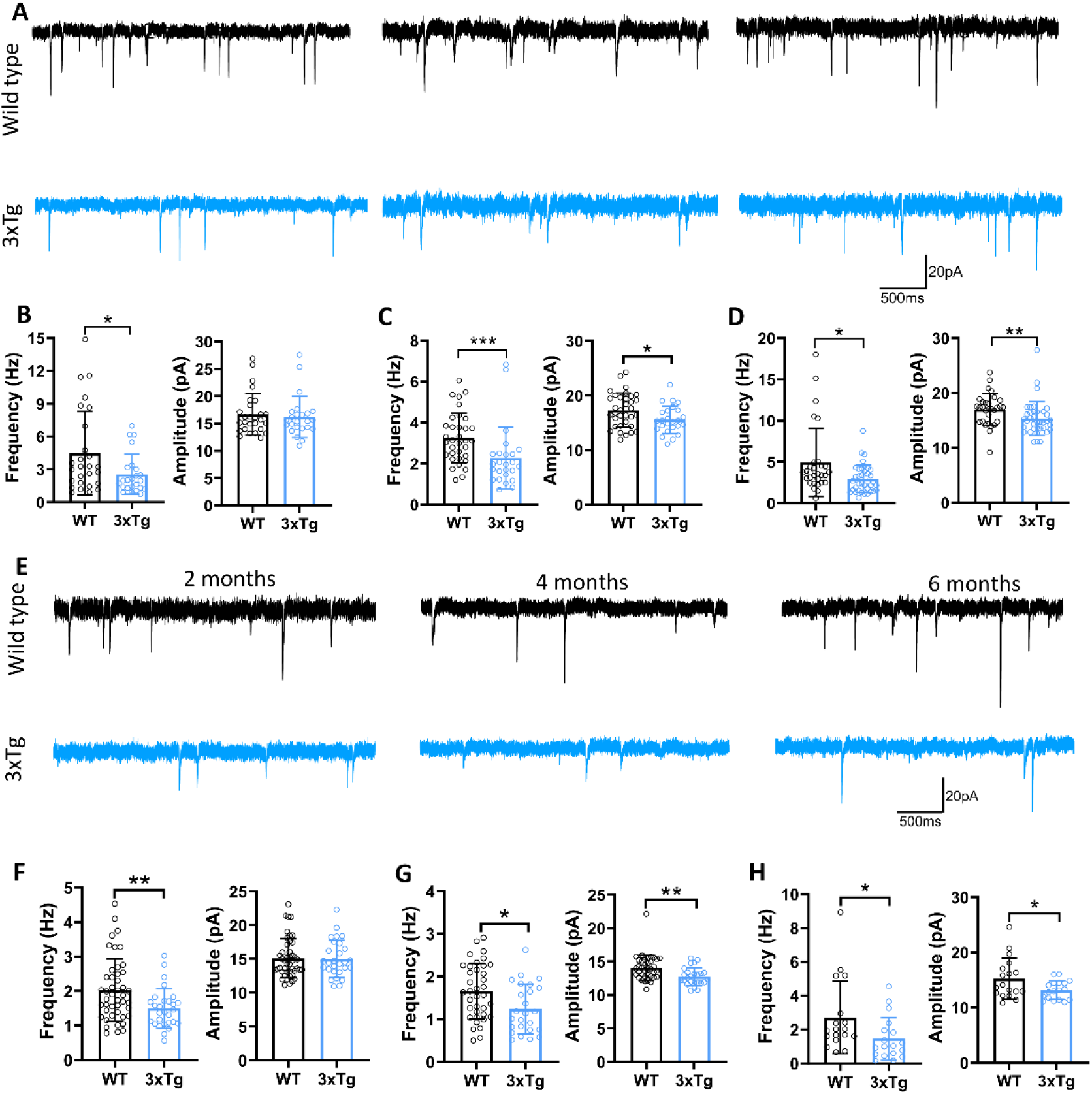
sEPSC and mEPSC in LC/NE neurons of young 3xTg mice. **A** Representative traces of spontaneous excitatory postsynaptic currents (sEPSCs) recorded from LC neurons at different ages. **B** sEPSC frequency and amplitude of LC neurons at 2 months of age. Left: sEPSC frequency (WT: 4.48 ± 3.83 Hz, n = 27 cells, N = 6 mice; 3xTg: 2.54 ± 1.82 Hz, n = 24 cells, N = 5 mice; Shapiro–Wilk tests, WT: W=0.819, p=0.0003, 3xTg: W=0.8, p=0.0003; Levene’s test, F_(1,9)_=2.351, p=0.160; unpaired two-tailed Mann-Whitney test, U=219p = 0.048). Right: sEPSC amplitude (WT: 16.67 ± 3.8 pA; 3xTg: 16.20 ± 3.78 pA; Shapiro – Wilk tests, WT: W=0.855, p=0.001, 3xTg: W=0.826, p=0.0008; Levene’s test, F_(1,9)_=0.808, p=0.392; unpaired two-tailed Mann-Whitney test, U=309p = 0.782). **C** sEPSC frequency and amplitude of LC neurons at 4 months of age. Left: sEPSC frequency (WT: 3.25 ± 1.22 Hz, n = 34 cells, N = 8 mice; 3xTg: 2.27 ± 1.5 Hz, n = 25 cells, N = 5 mice; Shapiro–Wilk tests, WT: W=0.969, p=0.445, 3xTg: W=0.738, p**<**0.001; Levene’s test, F_(1,11)_=1.228, p=0.291; unpaired two-tailed Mann-Whitney test, U=192p **<**0.001). Right: sEPSC amplitude (WT: 17.31 ± 3.13 pA; 3xTg: 15.57 ± 2.52 pA; Shapiro–Wilk tests, WT: W=0.972, p=0.525, 3xTg: W=0.975, p=0.77; Levene’s test, F_(1,11)_=0.033, p=0.86; unpaired two-tailed t test, t₅₇ = 2.274, p = 0.020). **D** sEPSC frequency and amplitude of LC neurons at 6 months of age. Left: sEPSC frequency (WT: 4.92 ± 4.11 Hz, n = 30 cells, N = 6 mice; 3xTg: 2.95 ± 1.7 Hz, n = 37 cells, N = 7 mice; Shapiro– Wilk tests, WT: W=0.717, p**<**0.001, 3xTg: W=0.899, p=0.003; Levene’s test, F_(1,11)_=2.29, p=0.158; unpaired two-tailed Mann-Whitney test, U=362p = 0.014). Right: sEPSC amplitude (WT: 17.01 ± 2.89 pA; 3xTg: 15.35 ± 3.15 pA; Shapiro–Wilk tests, WT: W=0.964, p=0.389, 3xTg: W=0.839, p**<**0.001; Levene’s test, F_(1,11)_=0.014, p=0.907; unpaired two-tailed Mann-Whitney test, U=314.5p = 0.002). **E** Representative traces of miniature excitatory postsynaptic currents (mEPSCs) recorded from LC neurons at different ages. **F** mEPSC frequency and amplitude of LC neurons at 2 months of age. Left: mEPSC frequency (WT: 2.03 ± 0.9 Hz, n = 45 cells, N = 10 mice; 3xTg: 1.50 ± 0.57 Hz, n = 28 cells, N = 7 mice; Shapiro–Wilk tests, WT: W=0.931, p=0.01, 3xTg: W=0.929, p=0.058; Levene’s test, F_(1,15)_=6.952, p=0.019; unpaired two-tailed Mann-Whitney test, U=402p = 0.009). Right: mEPSC amplitude (WT: 15.08 ± 2.91 pA; 3xTg: 15.01 ± 2.73 pA; Shapiro–Wilk tests, WT: W=0.906, p=0.001, 3xTg: W=0.948, p=0.178; Levene’s test, F_(1,15)_=1.408, p=0.254; unpaired two-tailed Mann-Whitney test, U=621.5, p = 0.926). **G** mEPSC frequency and amplitude of LC neurons at 4 months of age. Left: mEPSC frequency (WT: 1.65 ± 0.65 Hz, n = 37 cells, N = 7 mice; 3xTg: 1.24 ± 0.58 Hz, n = 25 cells, N = 5 mice; Shapiro– Wilk tests, WT: W=0.971, p=0.450, 3xTg: W=0.927, p=0.074; Levene’s test, F_(1,10)_=0.351, p=0.567; unpaired two-tailed t test, t₆₀ = 2.590, p = 0.010). Right: mEPSC amplitude (WT: 14.10 ± 1.89 pA; 3xTg: 12.76 ± 1.33 pA; Shapiro–Wilk tests, WT: W=0.837, p**<**0.001, 3xTg: W=0.952, p=0.272; Levene’s test, F_(1,10)_=1.909, p=0.197; unpaired two-tailed Mann-Whitney test, U=238.5p = 0.001). **H** mEPSC frequency and amplitude of LC neurons at 6 months of age. Left: mEPSC frequency (WT: 2.72 ± 2.14 Hz, n = 18 cells, N = 5 mice; 3xTg: 1.46 ± 1.26 Hz, n = 18 cells, N = 5 mice; Shapiro– Wilk tests, WT: W=0.786, p**<**0.001, 3xTg: W=0.861, p=0.013; Levene’s test, F_(1,8)_=1.251, p=0.296; unpaired two-tailed Mann-Whitney test, U=86p = 0.015). Right: mEPSC amplitude (WT: 15.26 ± 3.68 pA; 3xTg: 13.15 ± 1.67 pA; Shapiro–Wilk tests, WT: W=0.901, p**=**0.059, 3xTg: W=0.925, p=0.159; Levene’s test, F_(1,8)_=4.833, p=0.059; unpaired two-tailed t test, t₃₄ = 2.210, p = 0.030). Data are represented as mean ± SD. Each point represents an individual recorded cell. n, number of cells; N, number of animals. *p < 0.05; **p < 0.01.

sEPSC frequency compared to age-matched WT (Figure 3B). Importantly, sEPSC amplitude at this age was not significantly different between genotypes, indicating that the reduction in excitatory drive at 2 months is predominantly presynaptic in origin, reflecting decreased glutamate release from afferent terminals rather than alterations in postsynaptic glutamate receptor function. This finding suggests that the earliest detectable synaptic deficit in LC circuits involves suppression of excitatory neuron activity or reduced probability of neurotransmitter release from excitatory terminals, while the complement of postsynaptic AMPA and NMDA receptors remains intact.

The reduction in sEPSC frequency persisted at 4 months of age, with 3xTg mice continuing to display significantly decreased excitatory synaptic event rates compared to WT (Figure 3C). Notably, the statistical significance of this deficit increased relative to the 2-month time point, suggesting that the magnitude of the presynaptic deficit may be progressive or that the phenotype becomes more consistent across individual neurons and animals with advancing age. LC neurons from 4-month-old 3xTg mice exhibited not only persistently reduced sEPSC frequency but also significantly decreased sEPSC amplitude (Figure 3C). The emergence of reduced sEPSC amplitude at this age indicates the development of postsynaptic dysfunction, most likely reflecting a decrease in the number of functional AMPA receptors at glutamatergic synapses onto LC neurons, reduced single-channel conductance of these receptors, or both. As observed at 4 months, the synaptic phenotype in 3xTg mice at 6 months exhibited persistently reduced sEPSC frequency and significantly decreased sEPSC amplitude (Figure 3D), indicating both presynaptic and postsynaptic deficits. This temporal progression from purely presynaptic deficits at 2 months to combined presynaptic and postsynaptic impairments at 4 and 6 months, reveals a staged deterioration of excitatory synaptic transmission in the 3xTg model, with postsynaptic receptor dysfunction representing a later-emerging pathological feature that compounds the initial loss of presynaptic drive.

Next, to determine whether the observed reductions in sEPSC frequency reflect alterations in action potential-independent synaptic vesicle release or changes in the intrinsic properties of individual glutamatergic synapses, we recorded miniature EPSCs (mEPSCs) in the presence of the voltage-gated sodium channel blocker tetrodotoxin (TTX, 1 μM), which abolishes action potential-dependent neurotransmitter release and isolates spontaneous vesicle fusion events (Figure 3E). Under these conditions, mEPSC frequency provides a direct readout of the probability of spontaneous vesicle release (p) and/or the total number of functional excitatory synapses onto the recorded neuron, whereas mEPSC amplitude reflects the quantal size determined by the number and single-channel conductance of postsynaptic glutamate receptors(Kerchner and Nicoll, 2008; Prange and Murphy, 1999; Turrigiano, 2008).

At 2 months of age, analysis of mEPSCs revealed a significant reduction in event frequency in LC neurons from 3xTg mice compared to WT (Figure 3F). This finding is fully consistent with the reduced sEPSC frequency observed at the same age and demonstrates that the deficit in excitatory synaptic drive cannot be attributed solely to decreased action potential firing in presynaptic glutamatergic neurons. Instead, the reduced mEPSC frequency indicates presynaptic dysfunction at the level of individual synapses, which could manifest as: 1) decreased probability of spontaneous vesicle fusion at existing synapses, potentially resulting from alterations in the vesicle release machinery, reduced calcium sensitivity of the fusion apparatus, or changes in the readily releasable pool size; 2) a reduction in the total number of functional excitatory synapses onto LC neurons, reflecting either failed synapse formation, synapse loss, or conversion of previously active synapses to a functionally silent state; or 3) some combination of these mechanisms. Together, these findings demonstrate that diminished presynaptic excitatory drive underlies the reduction in excitatory synaptic transmission onto LC neurons. Importantly, mEPSC amplitude at 2 months was not significantly different between genotypes (Figure 3F), confirming that postsynaptic glutamate receptor complement remains normal at this early age, consistent with the sEPSC amplitude data.

Analysis of mEPSCs at 4 and 6 months of age (Figure 3G, H) revealed that the reduction in quantal event frequency persisted, indicating a sustained presynaptic deficit in glutamate release. In addition, mEPSC amplitudes were significantly decreased at both time points, consistent with the reduction observed in sEPSC amplitudes and pointing to emerging postsynaptic dysfunction. Notably, while the decrease in mEPSC frequency was already evident at 2 months and remained stable through 4 and 6 months, the reduction in mEPSC amplitude was not observed until 4 months, after which it persisted. These findings suggest a temporal progression in synaptic pathology, in which presynaptic impairments arise early and remain stable, followed by the onset of postsynaptic deficits that are subsequently maintained during early stages of AD.

### Enhanced GABAergic inhibitory transmission in LC of young 3xTg mice

The balance between excitatory and inhibitory synaptic drive is a fundamental determinant of neuronal excitability and network function, and disruption of this excitation-inhibition (E-I) balance has emerged as a critical pathophysiological mechanism in AD and other neurological disorders(Busche and Konnerth, 2016; Palop and Mucke, 2010). In the locus coeruleus, GABAergic inhibitory inputs arise from multiple sources, including local interneurons, afferents from the periaqueductal gray (PAG), and projections from other brainstem nuclei, and these inputs play important roles in regulating LC activity states, modulating phasic burst firing, and coordinating arousal and sleep-wake transition(Aston-Jones and Cohen, 2005; Breton-Provencher and Sur, 2019).

To comprehensively characterize inhibitory synaptic function in LC neurons during early AD progression, we performed voltage-clamp recordings of both spontaneous inhibitory postsynaptic currents (sIPSCs) (Figure 4A) and miniature inhibitory postsynaptic currents (mIPSCs) (Figure 4E) in 3xTg and wild-type mice at 2, 4, and 6 months of age. As in our analysis of excitatory transmission, the combined assessment of sIPSCs and mIPSCs provides complementary insights. sIPSCs reflect the overall inhibitory synaptic input, encompassing both action potential–dependent and – independent GABA release, whereas mIPSCs, recorded in the presence of tetrodotoxin (TTX), isolate spontaneous vesicular fusion events and thus offer direct information on quantal GABA release and the functional status of individual inhibitory synapses.

**Figure 4.**
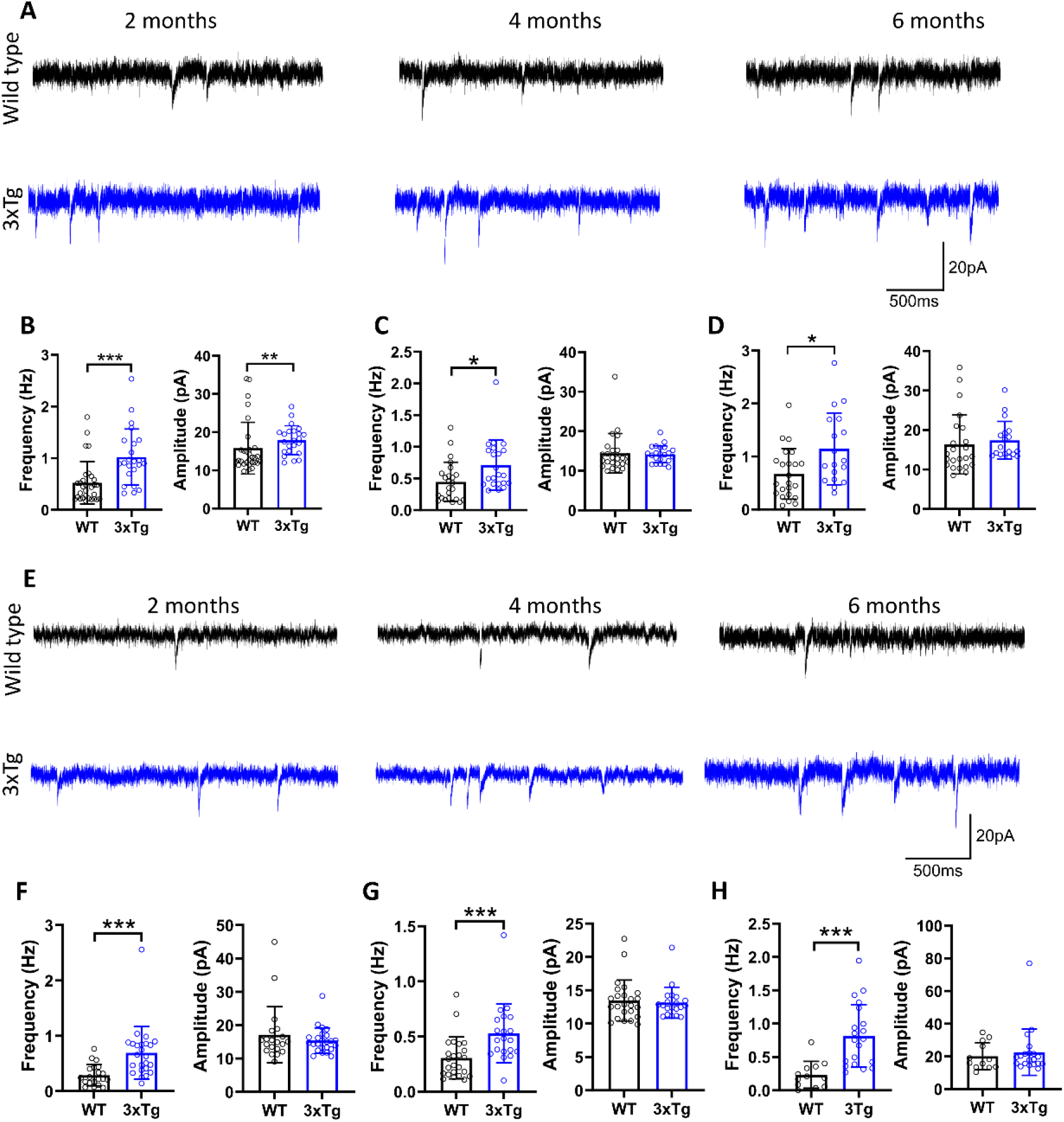
sIPSC and mIPSC in LC/NE neurons of young 3xTg mice. **A** Representative traces of spontaneous inhibitory postsynaptic currents (sIPSCs) recorded from LC neurons at different ages. **B** sIPSC frequency and amplitude of LC neurons at 2 months of age. Left: sIPSC frequency (WT: 0.52 ± 0.41 Hz, n = 30 cells, N = 5 mice; 3xTg: 1.023 ± 0.542 Hz, n = 23 cells, N = 5 mice; Shapiro–Wilk tests, WT: W=0.762, p**<**0.001, 3xTg: W=0.917, p=0.058; Levene’s test, F_(1,8)_=1.865, p=0.209; unpaired two-tailed Mann-Whitney test, U=132p **<**0.001). Right: sIPSC amplitude (WT: 15.80 ± 6.77 pA; 3xTg: 17.902 ± 3.782 pA; Shapiro–Wilk tests, WT: W=0.713, p**<**0.001, 3xTg: W=0.971, p=0.709; Levene’s test, F_(1,8)_=1.04, p=0.338; unpaired two-tailed Mann-Whitney test, U=184p = 0.003). **C** sIPSC frequency and amplitude of LC neurons at 4 months of age. Left: sIPSC frequency (WT: 0.45 ± 0.31 Hz, n = 24 cells, N = 4 mice; 3xTg: 0.71 ± 0.4 Hz, n = 22 cells, N = 4 mice; Shapiro–Wilk tests, WT: W=0.878, p**=**0.008, 3xTg: W=0.811, p**<**0.001; Levene’s test, F_(1,6)_=1.095, p=0.336; unpaired two-tailed Mann-Whitney test, U=154p = 0.015). Right: sIPSC amplitude (WT: 14.40 ± 5.0 pA; 3xTg: 14.16 ± 2.14 pA; Shapiro – Wilk tests, WT: W=0.709, p**<**0.001, 3xTg: W=0.955, p**=**0.396; Levene’s test, F_(1,6)_=3.764, p=0.100; unpaired two-tailed Mann-Whitney test, U=221.5p = 0.356). **D** sIPSC frequency and amplitude of LC neurons at 6 months of age. Left: sIPSC frequency (WT: 0.67 ± 0.48 Hz, n = 23 cells, N = 5 mice; 3xTg: 1.144 ± 0.678 Hz, n = 18cells, N = 5 mice; Shapiro–Wilk tests, WT: W=0.919, p**=**0.065, 3xTg: W=0.917, p**=**0.114; Levene’s test, F_(1,8)_=2.837, p=0.131; unpaired two-tailed t test, t_36_ = 2.632, p=0.012). Right: sIPSC amplitude (WT: 16.89 ± 3.19 pA; 3xTg: 17.391 ± 4.776 pA; Shapiro–Wilk tests, WT: W=0.828, p**=**0.001, 3xTg: W=0.821, p**=**0.003; Levene’s test, F_(1,8)_=0.147, p=0.712; unpaired two-tailed Mann-Whitney test, U=143p = 0.096). **E** Representative traces of miniature inhibitory postsynaptic currents (mIPSCs) recorded from LC neurons at different ages. **F** mIPSC frequency and amplitude of LC neurons at 2 months of age. Left: mIPSC frequency (WT: 0.29 ± 0.19 Hz, n = 20 cells, N = 5 mice; 3xTg: 0.695 ± 0.474 Hz, n = 24 cells, N = 5 mice; Shapiro– Wilk tests, WT: W=0.915, p**=**0.079, 3xTg: W=0.734, p**<**0.001; Levene’s test, F_(1,8)_=1.892, p=0.206; unpaired two-tailed Mann-Whitney test,U=72, p < 0.001). Right: mIPSC amplitude (WT: 17.13 ± 8.39 pA; 3xTg: 15.387 ± 3.819 pA; Shapiro–Wilk tests, WT: W=0.689, p**<**0.001, 3xTg: W=0.847, p**=**0.002; Levene’s test, F_(1,8)_=4.942, p=0.057; unpaired two-tailed Mann-Whitney test, U=232p = 0.856). **G** mIPSC frequency and amplitude of LC neurons at 4 months of age. Left: mIPSC frequency (WT: 0.31 ± 0.19 Hz, n = 24 cells, N = 5 mice; 3xTg: 0.529 ± 0.268 Hz, n = 20 cells, N = 5 mice; Shapiro– Wilk tests, WT: W=0.848, p**=**0.002, 3xTg: W=0.836, p**=**0.003; Levene’s test, F_(1,8)_=0.821, p=0.391; unpaired two-tailed Mann-Whitney test, U=97p**<**0.001). Right: mIPSC amplitude (WT: 13.47 ± 3.06 pA; 3xTg: 13.148 ± 2.316 pA; Shapiro–Wilk tests, WT: W=0.856, p**=**0.003, 3xTg: W=0.74, p**<**0.001; Levene’s test, F_(1,8)_=0.016, p=0.901; unpaired two-tailed Mann-Whitney test, U=231.5p = 0.847). **H** mIPSC frequency and amplitude of LC neurons at 6 months of age. Left: mIPSC frequency (WT: 0.23 ± 0.2 Hz, n = 12 cells, N = 4 mice; 3xTg: 0.817 ± 0.469 Hz, n = 20 cells, N = 5 mice; Shapiro– Wilk tests, WT: W=0.900, p**=**0.159, 3xTg: W=0.915, p**=**0.079; Levene’s test, F_(1,7)_=0.912, p=0.371; unpaired two-tailed t test, t₂₄ = 4.168, p < 0.001). Right: mIPSC amplitude (WT: 20.16 ± 8.28 pA; 3xTg: 22.636 ± 14.196 pA; Shapiro–Wilk tests, WT: W=0.881, p**=**0.089, 3xTg: W=0.596, p**<**0.001; Levene’s test, F_(1,7)_=0.002, p=0.968; unpaired two-tailed Mann-Whitney test, U=104p = 0.552). Data are represented as mean ± SD. Each point represents an individual recorded cell. n, number of cells; N, number of animals. *p < 0.05; **p < 0.01; ***p < 0.001.

Spontaneous IPSCs were recorded from LC neurons voltage-clamped at −60 mV in the presence of the AMPA receptor antagonist CNQX (10 μM) and the NMDA receptor antagonist D-AP5 (50 μM) to block all glutamatergic transmission and isolate GABAergic currents. In striking contrast to the reduced excitatory synaptic activity observed in 3xTg mice, our analysis revealed robust and persistent enhancement of inhibitory synaptic transmission across all three ages examined, indicating a dramatic shift in the excitation-inhibition balance toward increased inhibitory drive (Figure 4A). At 2 months of age, LC neurons from 3xTg mice displayed a striking increase in sIPSC frequency compared to age-matched WT mice (Figure 4B). The magnitude of this effect was substantial, achieving high statistical significance, and indicating that LC neurons in 2-month-old 3xTg mice experience dramatically increased GABAergic inhibitory bombardment.

The enhancement of sIPSC frequency at 2 months represents a maximal upregulation of inhibitory synaptic strength, involving increases in presynaptic GABA release or number of functional GABAergic synapse. This stands in stark contrast to the presynaptic deficit observed for excitatory transmission at the same age (reduced sEPSC frequency with normal amplitude) and suggests that excitatory and inhibitory synaptic systems are differentially regulated in the 3xTg model, with inhibitory synapses showing a gain-of-function phenotype while excitatory synapses exhibit loss of function. At 4 months of age, the enhancement of sIPSC frequency persisted in 3xTg mice, with LC neurons continuing to display significantly elevated inhibitory event rates compared to WT mice (Figure 4C). By 6 months of age, the enhancement of sIPSC frequency in 3xTg mice was even more pronounced, with the statistical significance increasing relative to earlier ages (Figure 4D). This progressive intensification of the frequency phenotype suggests that the mechanisms driving enhanced inhibitory transmission become increasingly robust or widespread with advancing age and disease progression. The p values across ages (0.003 at 2 months → 0.01 at 4 months → 0.0009 at 6 months) show that while there is some variability, the overall effect remains highly significant and persistent. Importantly, sIPSC amplitudes at all ages were not significantly different between genotypes, confirming that the primary locus of inhibitory synaptic enhancement in older 3xTg mice is presynaptic, involving increased GABA release rather than enhanced postsynaptic receptor function.

To determine whether the observed increases in sIPSC frequency reflect alterations in action potential-independent vesicular GABA release or changes in the intrinsic properties of individual inhibitory synapses, we recorded miniature IPSCs (mIPSCs) in the presence of TTX (1 μM) to block action potential-dependent neurotransmitter release (Figure 4E). mIPSCs were recorded at −60 mV using potassium-based internal solution to maintain physiological chloride gradients and ensure that GABA*_A_* receptor activation produces inward currents. Under these conditions, mIPSC frequency directly reflects the probability of spontaneous GABA vesicle fusion and/or the total number of functional inhibitory synapses onto the recorded neuron, while mIPSC amplitude indicates the quantal size determined by postsynaptic GABA*_A_* receptor number and single-channel conductance. At 2 months of age, analysis of mIPSCs revealed a highly significant increase in event frequency in LC neurons from 3xTg mice compared to WT mice (Figure 4F). This finding is fully consistent with the elevated sIPSC frequency observed at the same age and demonstrates that the enhanced inhibitory synaptic drive cannot be attributed solely to increased action potential firing in presynaptic GABAergic neurons. Instead, the elevated mIPSC frequency indicates intrinsic changes at the level of individual inhibitory synapses, which could manifest as: 1) increased probability of spontaneous GABA vesicle fusion at existing synapses, potentially resulting from enhanced calcium sensitivity of the release machinery, increased size of the readily releasable vesicle pool, or alterations in vesicle priming proteins; 2) an increase in the total number of functional inhibitory synapses onto LC neurons, reflecting enhanced synapse formation, stabilization of nascent synapses, or conversion of previously silent synapses to functionally active states; or 3) some combination of these mechanisms. Critically, mIPSC amplitude at 2 months was not significantly different between genotypes (Figure 4F), the normal mIPSC amplitude indicates that the baseline quantal GABA*_A_* receptor function at individual synapses is intact, with the enhancement primarily reflecting presynaptic mechanisms.

The elevation in mIPSC frequency persisted at 4 months of age, with 3xTg neurons continuing to display significantly increased inhibitory event rates compared to WT mice (Figure 4G). The sustained enhancement of mIPSC frequency across the 2–4-month period indicates that the presynaptic mechanisms driving increased quantal GABA release are established early and maintained throughout this developmental window. Consistent with the pattern observed at 2 months, mIPSC amplitude at 4 months was not significantly different between genotypes, confirming that postsynaptic GABA*_A_* receptor function remains normal and that the primary locus of inhibitory synaptic enhancement is presynaptic. The presynaptic enhancement of inhibitory transmission persisted through 6 months of age, with 3xTg mice displaying continued elevation in mIPSC frequency (Figure 4H). As observed at earlier ages, mIPSC amplitude at 6 months was not significantly different between genotypes, indicating that the normal quantal GABA*_A_* receptor function is maintained throughout the early disease period despite sustained enhancement of presynaptic GABA release.

### Voltage-gated K⁺ currents are reduced in LC/NE neurons of young 3xTg mice

To determine whether altered voltage-gated K⁺ conductance contributes to the abnormal firing properties of LC noradrenergic neurons in young 3xTg mice, we recorded outward K⁺ currents in response to a series of depolarizing voltage steps. Both WT and 3xTg LC neurons displayed rapidly activating outward currents followed by sustained current components during prolonged depolarization (Fig 5A). Quantification of the current–voltage relationships revealed a significant reduction in peak K⁺ current density in 3xTg neurons compared with WT controls, particularly at strongly depolarized membrane potentials (Fig 5B). Steady-state K⁺ current density measured near the end of each voltage step was also significantly reduced in 3xTg neurons (Fig 5C). Tail-current density showed a similar overall reduction in 3xTg neurons across depolarizing test potentials (Fig 5D).

**Figure 5.**
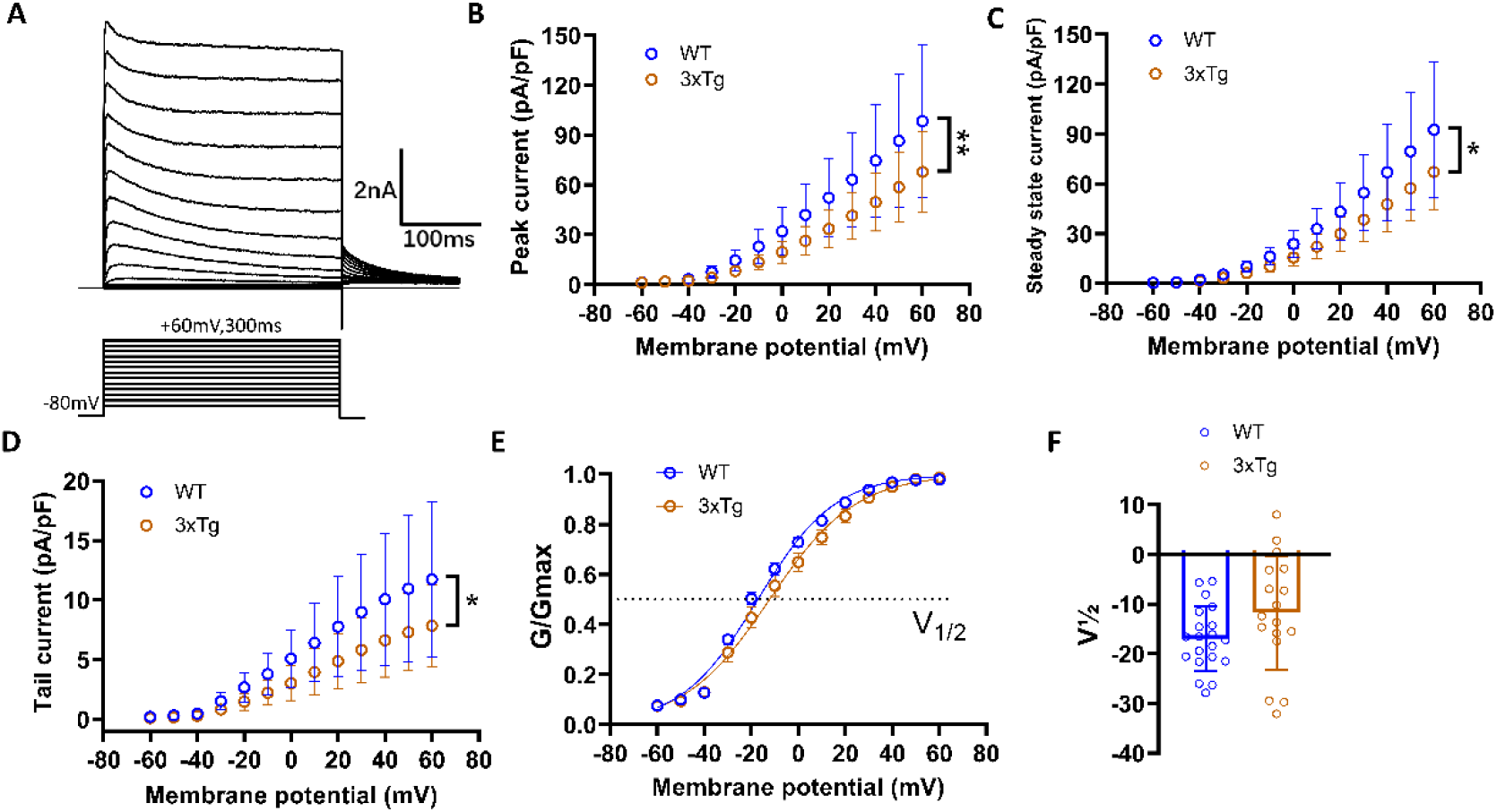
Voltage-gated K⁺ current density is reduced in LC/NE neurons of young 3xTg mice. **A** Representative traces of total outward K⁺ currents recorded from LC neurons during depolarizing voltage steps from −60 mV to +60mV (WT: n=20 cells, N=5 mice; 3xTg: n=17 cells, N=5mice). Peak, steady-state, and tail currents were measured as indicated. **B** Current density–voltage (I–V) relationship of peak outward K⁺ currents. Peak current amplitudes were normalized to membrane capacitance (pA/pF). LC neurons from 3xTg mice exhibited significantly reduced peak K⁺ current density compared with WT neurons (two-way repeated-measures ANOVA, P = 0.007, F_1,35_=8.359). **C** Current density–voltage relationship of steady-state outward K⁺ currents measured at the end of each depolarizing voltage step. Steady-state K⁺ current density was significantly reduced in 3xTg neurons compared with WT neurons (two-way repeated-measures ANOVA, P = 0.011, F_1,35_=7.189). **D** Tail current density measured immediately after repolarization following depolarizing voltage steps. Tail current density was significantly decreased in 3xTg LC neurons relative to WT controls (two-way repeated-measures ANOVA, P = 0.015, F_1,35_=6.531). **E** Voltage dependence of K⁺ channel activation. Conductance (G) was calculated from tail currents, normalized to maximal conductance (G/G_max), and fitted with a Boltzmann function to determine the half-activation voltage (V₁/₂). Activation curves were largely similar between WT and 3xTg neurons. **F** Summary of the half-activation voltage (V₁/₂) derived from Boltzmann fits. No statistically significant difference in V₁/₂ was detected between WT and 3xTg LC neurons (WT: −16.959±6.452 mV, n=20 cells, N=5 mice; 3xTg: −11.771±11.399 mV, n=17 cells, N=5 mice; Shapiro–Wilk tests, WT: W=0.966, p**=**0.669, 3xTg: W=0.958, p**=**0.602; Levene’s test, F_(1,8)_=21.304, p=0.002; unpaired two-tailed Welch’s *t* test, p = 0.109), indicating that the voltage dependence of K⁺ channel activation was largely preserved despite reduced K⁺ current density. Data are presented as mean ± SD. In B– D, symbols and error bars represent the mean ± SD at each membrane potential; individual observations are not displayed at every voltage step to preserve curve clarity. In E, conductance–voltage relationships were fitted using a Boltzmann function. In F, bars represent mean ± SD, and individual data points are presented. Sample sizes are reported as n = cells/N = mice. *P < 0.05; **P < 0.01.

To determine whether the reduced K⁺ currents reflected altered channel gating, conductance was normalized to the maximal conductance and plotted as a function of membrane potential. Boltzmann analysis showed largely overlapping activation curves between WT and 3xTg neurons (Fig 5E). Consistent with this observation, the half-maximal activation voltage (V₁/₂) was not significantly different between groups (Fig 5F). Thus, the reduction in K⁺ current was not accompanied by a detectable shift in its voltage dependence of activation. These findings demonstrate an early reduction in voltage-gated K⁺ conductance in LC noradrenergic neurons of 3xTg mice. Because K⁺ currents normally oppose membrane depolarization and promote action-potential repolarization, their reduction may contribute to elevated spontaneous activity, although it does not account for the impaired firing response to sustained depolarizing input.

### Voltage-gated Na⁺ current density is reduced in LC/NE neurons of young 3xTg mice

We next examined whether voltage-gated Na⁺-channel dysfunction contributes to the reduced evoked firing capacity of LC noradrenergic neurons in young 3xTg mice. Voltage-gated Na⁺ currents were elicited by depolarizing voltage steps from a hyperpolarized holding potential. Representative current traces showed rapidly activating and inactivating inward currents in neurons from both WT and 3xTg mice (Fig 6C). However, analysis of the current–voltage relationship revealed that Na⁺ current density was significantly reduced in 3xTg LC neurons across a range of activating membrane potentials (Fig 6D).

**Figure 6.**
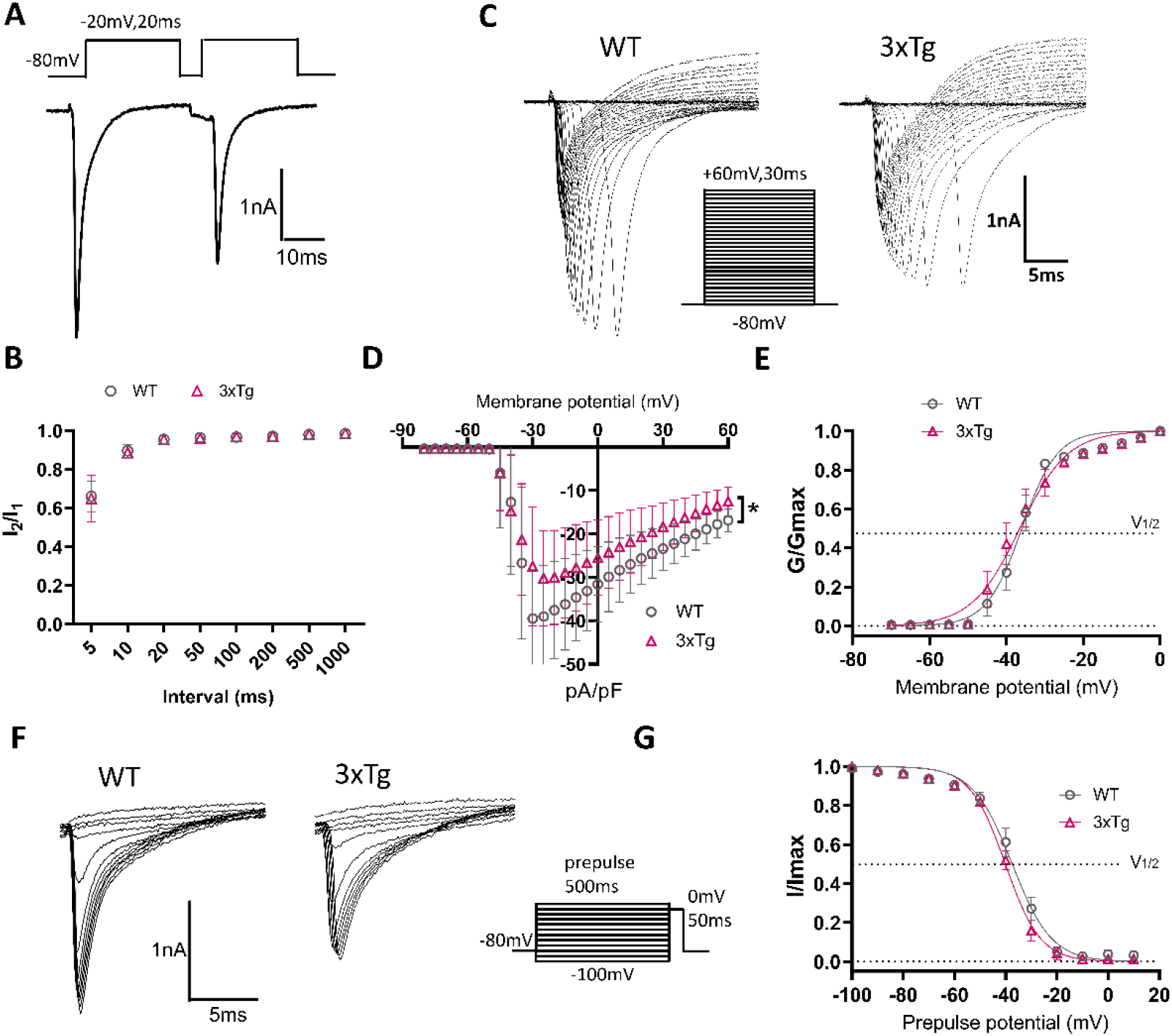
Voltage-gated Na⁺ conductance is reduced in LC/NE neurons of young 3xTg mice. **A** Representative voltage-clamp recording showing the protocol used to assess recovery from inactivation of voltage-gated Na⁺ channels. Two identical depolarizing test pulses were separated by recovery intervals ranging from 5 to 1000 ms. **B** Recovery from inactivation of Na⁺ currents was quantified as the ratio of the second to the first peak current (I₂/I₁). Recovery kinetics were comparable between WT and 3xTg LC neurons across all recovery intervals, indicating no significant change in recovery from inactivation (WT: n=17 cells, N=5 mice; 3xTg: n=15 cells, N=5 mice; two way repeated measure ANOVA, F_(1,30)_=0.186, p=0.67). **C** Representative traces of voltage gated Na⁺ currents recorded during depolarizing voltage steps. **D** Current density–voltage (I–V) relationship of voltage-gated Na⁺ currents. Peak current amplitudes were normalized to membrane capacitance (pA/pF). 3xTg neurons exhibited significantly reduced Na⁺ current density compared with WT neurons (WT: n=16 cells, N=5 mice; 3xTg: n=13 cells, N=4 mice; two-way repeated-measures ANOVA, p = 0.018, F_(1,27)_=6.339). **E** Voltage dependence of Na⁺ channel activation. Conductance was calculated from peak Na⁺ currents and normalized to the maximal conductance (G/G_max). Data were fitted with a Boltzmann function to determine the half-activation voltage (V₁/₂). No significant difference in activation properties was observed between WT and 3xTg neurons. **F** Representative Na⁺ current traces obtained using the steady-state inactivation protocol following conditioning prepulses at different membrane potentials. **G** Steady-state inactivation curves of voltage-gated Na⁺ channels. Peak currents were normalized to the maximal current (I/I_max) and fitted with a Boltzmann function. The voltage dependence of steady-state inactivation was similar between WT and 3xTg neurons, with no significant change in V₁/₂.. In B, D, E, and G, symbols represent group means and error bars represent SD. Individual observations are not displayed at every interval or voltage step to preserve the clarity of the recovery, current – voltage, activation, and inactivation curves. Sample sizes are reported as n = cells/N = mice for each condition. *p < 0.05.

To determine whether this reduction was associated with altered voltage-dependent gating, normalized conductance was plotted against membrane potential and fitted with a Boltzmann function. The voltage dependence of Na⁺-channel activation was comparable between WT and 3xTg neurons, with substantially overlapping activation curves and no detectable shift in the half-maximal activation voltage (Fig 6E). Steady-state inactivation curves were also similar between groups, indicating that the voltage dependence of channel availability was preserved in 3xTg neurons (Fig 6F–G).

We further assessed recovery from inactivation using a paired-pulse protocol with variable recovery intervals. The recovery time course was comparable between WT and 3xTg LC neurons, with no significant group difference across the tested intervals (Fig 6A–B). Therefore, the reduction in Na⁺ current density was not explained by altered activation, steady-state inactivation, or recovery from inactivation. Collectively, these findings indicate a reduction in functional voltage-gated Na⁺ conductance rather than a major alteration in channel-gating kinetics. Reduced Na⁺ current availability would limit action-potential initiation and repetitive spike generation during sustained depolarization, providing a plausible ionic mechanism for the impaired input-driven firing observed in 3xTg LC/NE neurons.

### Altered excitatory and inhibitory synaptic organization in LC of young 3xTg mice

To determine whether the functional synaptic imbalance observed in LC was accompanied by structural alterations in local synaptic inputs, we quantified presynaptic and postsynaptic puncta surrounding TH-positive LC neurons. Putative excitatory synapses were identified by the spatially colocalized or closely opposed presynaptic glutamatergic marker VGLUT2 and postsynaptic scaffold protein PSD95 (Figure 7A). Compared with WT controls, 3xTg mice exhibited significantly lower densities of both VGLUT2-positive and PSD95-positive puncta in LC. Consistent with these reductions, the density of paired VGLUT2/PSD95 puncta was also significantly decreased in 3xTg mice, indicating a loss or reduced abundance of putative excitatory synaptic contacts (Figure 7B).

**Figure 7.**
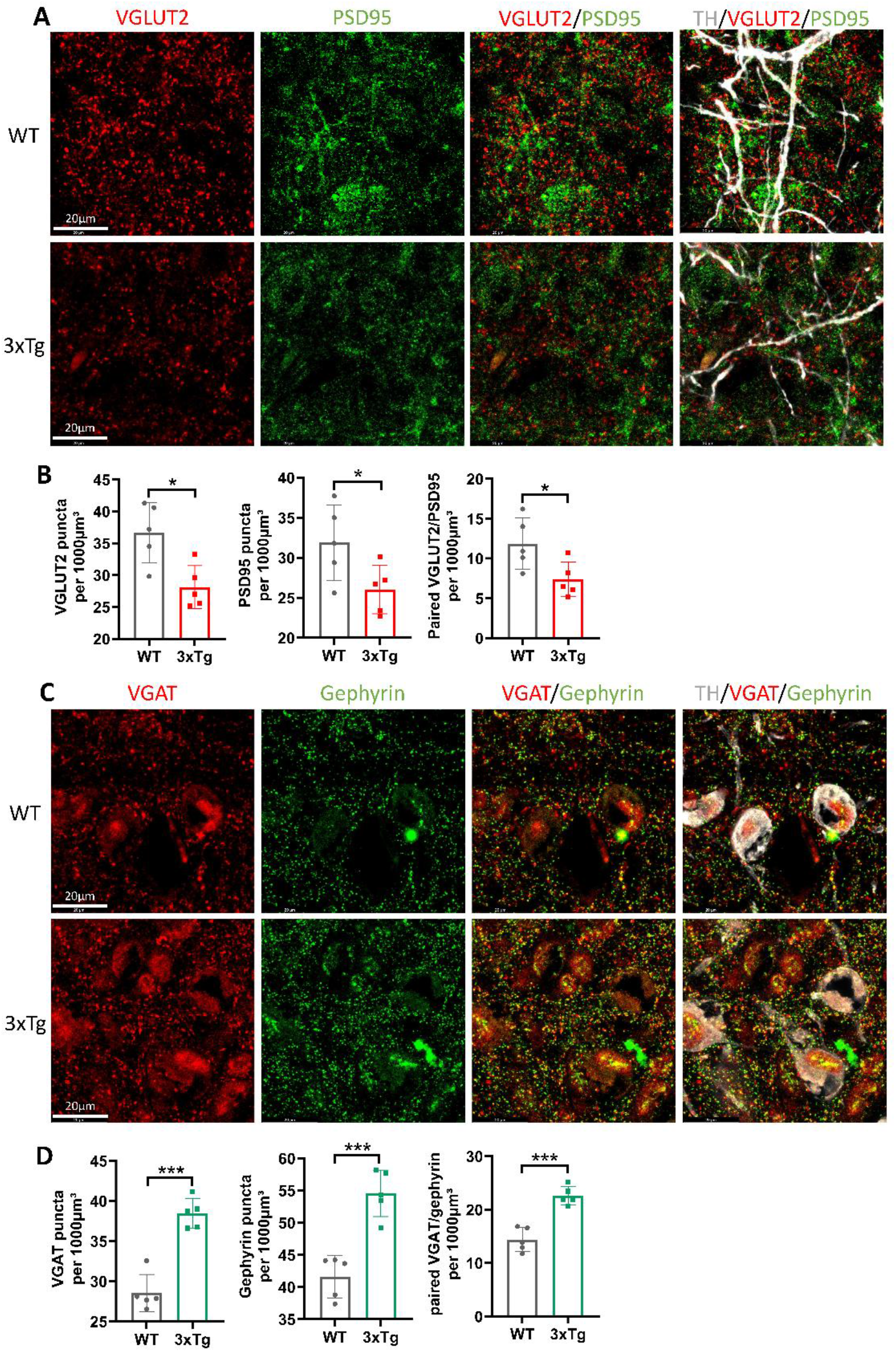
Reduced excitatory and increased inhibitory synaptic marker density in LC of young 3xTg mice. **A** Representative confocal images of VGLUT2 (red), PSD95 (green), and TH (white) immunoreactivity in the LC of WT and 3xTg mice. Merged images show the spatial distribution of VGLUT2 and PSD95 puncta and their relationship to TH-positive LC/NE neurons. **B** Quantification of VGLUT2-positive puncta, PSD95-positive puncta, and paired VGLUT2–PSD95 puncta per 1,000 µm³. The densities of VGLUT2-positive puncta (WT=36.68±4.72, N=5 mice; 3xTg=28.14±3.36, N=5 mice; Shapiro–Wilk tests, WT: W=0.933, p**=**0.616, 3xTg: W=0.893, p**=**0.371; Levene’s test, F_(1,8)_=0.569, p=0.472; unpaired two-tailed t test: *t*8=3.297, p=0.01), PSD95-positive puncta (WT=31.9 ± 4.72; 3xTg=26.02 ± 3.02; Shapiro – Wilk tests, WT: W=0.989, p**=**0.977, 3xTg: W=0.936, p**=**0.637; Levene’s test, F_(1,8)_=0.825, p=0.390; unpaired two-tailed t test: *t*8=2.346, p=0.04), and paired VGLUT2–PSD95 objects (WT=11.86±3.22; 3xTg=7.34 ± 2.17; Shapiro – Wilk tests, WT: W=0.962, p**=**0.824, 3xTg: W=0.919, p**=**0.524; Levene’s test, F_(1,8)_=1.299, p=0.287; unpaired two-tailed t test: *t*8=2.6, p=0.03) were significantly reduced in 3xTg mice compared with WT controls. **C** Representative confocal images of VGAT (red), gephyrin (green), and TH (white) immunoreactivity in the LC of WT and 3xTg mice. Merged images illustrate the distribution of inhibitory presynaptic and postsynaptic markers surrounding TH-positive LC neurons. **D** Quantification of VGAT-positive puncta, gephyrin-positive puncta, and paired VGAT– gephyrin puncta per 1,000 µm³. The densities of VGAT-positive puncta (WT=28.55± 2.32, N=5 mice; 3xTg=38.46±1.88, N=5 mice; Shapiro–Wilk tests, WT: W=0.783, p**=**0.058, 3xTg: W=0.907, p**=**0.448; Levene’s test, F_(1,8)_=0.048, p=0.832; unpaired two-tailed t test: *t*8=7.42, p<0.001), gephyrin-positive puncta (WT=41.63 ± 3.28; 3xTg=54.57 ± 3.62; Shapiro– Wilk tests, WT: W=0.788, p**=**0.065, 3xTg: W=0.925, p**=**0.561; Levene’s test, F_(1,8)_=0.027, p=0.874; unpaired two-tailed t test: *t*8=5.924, p<0.001), and paired VGAT–gephyrin objects (WT=14.38±2.25; 3xTg=22.59±1.75; Shapiro–Wilk tests, WT: W=0.902, p**=**0.424, 3xTg: W=0.952, p**=**0.753; Levene’s test, F_(1,8)_=0.797, p=0.398; unpaired two-tailed t test: *t*8=6.444, p<0.001) were significantly increased in 3xTg mice relative to WT controls. Data are presented as mean ± SD; each point represents one mouse. *p< 0.05, *** p< 0.001, scale bars: 20 µm.

In contrast, analysis of inhibitory synaptic markers revealed the opposite pattern. The density of VGAT-positive presynaptic puncta was markedly increased in LC of 3xTg mice relative to WT controls. Gephyrin-positive postsynaptic puncta were similarly increased. Consequently, the density of paired VGAT/gephyrin puncta, representing putative inhibitory synaptic contacts, was significantly elevated in 3xTg mice (Figure 7C, D).

Together, these findings reveal a bidirectional reorganization of synaptic inputs in LC of young 3xTg mice, characterized by reduced excitatory and increased inhibitory synaptic marker density. These anatomical changes are consistent with the electrophysiological evidence of diminished excitatory and enhanced inhibitory synaptic transmission onto LC/NE neurons, supporting the presence of an early shift in the local excitation–inhibition balance before substantial LC neuronal loss.

## Discussion

In AD, LC is among the earliest regions to accumulate tau pathology and degenerate (Beardmore et al., 2021; Braak and Del Tredici, 2011; Bueichekú et al., 2024; Hary et al., 2025; Matchett et al., 2021), and its dysfunction is linked to early neuropsychiatric symptoms and cognitive decline (Bouret and Sara, 2005; Feinstein et al., 2016; Giorgi et al., 2020; Kazim et al., 2017; Mercan and Heneka, 2022; Reyes, 2025; Robertson, 2013; Sara, 2009). However, whether functional impairment precedes overt LC degeneration, and what form this impairment takes, has remained unclear. To address this, we used whole-cell electrophysiology and quantitative immunofluorescence to characterize synaptic transmission, intrinsic excitability, and voltage-gated currents in LC neurons from young 3xTg mice, prior to substantial neurodegeneration. We identified pronounced excitation-inhibition imbalance, with reduced excitatory synaptic input and enhanced inhibitory transmission, together with a striking dissociation between elevated spontaneous pacemaking and impaired evoked firing. These findings indicate that synaptic and intrinsic dysfunction emerge early within the noradrenergic system and may represent a critical initiating event in AD pathophysiology, extending the well-established principle that synaptic dysfunction precedes neuronal loss and cognitive decline in AD to the LC(Scheff et al., 2007; Selkoe, 2002; Terry et al., 1991). To our knowledge, this represents the first direct electrophysiological characterization of synaptic transmission dynamics in LC neurons during prodromal AD. While previous studies have documented tau pathology and eventual LC degeneration in AD(Beardmore et al., 2021; Braak and Del Tredici, 2011; Bueichekú et al., 2024; Hary et al., 2025; Matchett et al., 2021), the functional consequences at the synaptic level have remained unexplored. Our demonstration of bidirectional synaptic changes including simultaneous reduction in excitation and enhancement of inhibition reveals a more complex dysregulation than previously appreciated and suggests that LC pathophysiology involves active synaptic remodeling rather than simple synaptic loss.

Several factors may help explain the unexpected dissociation we observed between spontaneous and evoked firing. Despite reduced intrinsic excitability, LC neurons exhibited elevated spontaneous firing rates, revealing an uncoupling of these two modes of activity that has been observed in other pathological contexts and may reflect compensatory or maladaptive homeostatic responses to declining excitatory input(Turrigiano, 2011). In the LC, spontaneous firing maintains baseline arousal and network tone(Aston-Jones and Cohen, 2005; Bateman et al., 2012; Caestecker et al., 2025; Carter et al., 2010; Grimm et al., 2024); increased baseline firing combined with impaired evoked responsiveness may therefore compromise the dynamic regulation of brain states needed for appropriate behavioral responses. This paradoxical hyperactivity despite diminished responsiveness carries direct implications for early AD symptomatology: an LC unable to appropriately modulate firing in response to physiological demand could contribute to the attention deficits, sleep disruption, and cognitive fluctuations observed in early-stage AD patients(Bouret and Sara, 2005; Feinstein et al., 2016; Giorgi et al., 2020; Reyes, 2025). Moreover, aberrant noradrenergic output from dysfunctional LC neurons may propagate network instability to downstream regions, potentially accelerating pathology spread throughout cortical and hippocampal circuits.

Our findings also bear on the relationship between LC functional dysregulation and degeneration. The LC is among the earliest regions to accumulate tau pathology in AD (Beardmore et al., 2021; Braak and Del Tredici, 2011; Bueichekú et al., 2024; Hary et al., 2025; Matchett et al., 2021), and degeneration of its projections correlates with cognitive impairment and network instability (Kazim et al., 2017; Mercan and Heneka, 2022; Robertson, 2013; Sara, 2009). By showing that functional dysregulation precedes substantial neurodegeneration, our results support a model in which early synaptic imbalance alters noradrenergic output and amplifies the vulnerability of downstream cortical and hippocampal circuits, challenging the traditional view that tau accumulation alone drives LC pathology. Instead, profound functional alterations appear to occur even before substantial cell loss, positioning synaptic dysfunction as a potential driver, rather than merely a consequence, of proteinopathy. This temporal sequence suggests a therapeutic window during which interventions targeting synaptic stability could prevent or delay the cascade of downstream pathology.

These implications likely extend beyond the LC itself. As the brain’s primary source of norepinephrine, the LC modulates synaptic plasticity, neuroinflammation, and metabolic function across widespread brain regions(Aston-Jones and Cohen, 2005; Caestecker et al., 2025; Carter et al., 2010; Grimm et al., 2024; Schwarz and Luo, 2015); early LC dysfunction could therefore act as a catalyst that destabilizes multiple neural networks simultaneously, offering one explanation for the multifaceted nature of early AD symptoms. Our identification of specific synaptic and ionic alterations further provides concrete candidate targets for intervention: strategies aimed at restoring excitatory-inhibitory balance or normalizing LC firing patterns during prodromal stages could potentially halt or reverse early network dysfunction before irreversible neuronal loss occurs.

Taken together, these results indicate that LC dysfunction in AD begins with functional synaptic and ionic disruption rather than with cellular degeneration, reframing how early AD pathology may emerge in this system and providing a mechanistic foundation for interventions aimed at preventing or delaying disease progression before it becomes irreversible.

## Limitations and future perspectives

Several methodological considerations should be taken into account when interpreting these findings. This study used the 3xTg mouse model, which, while recapitulating key amyloid and tau features of AD, may not fully capture the complexity of human disease; validation in additional models and, ultimately, human tissue will be important. Electrophysiological recordings were performed in acute brain slices, which lack the complete network connectivity and neuromodulatory inputs present in vivo; future in vivo recordings will be needed to confirm whether the synaptic alterations observed here translate into comparable functional changes in behaving animals. Although we identified a pronounced excitation-inhibition imbalance, we did not characterize the specific sources of excitatory and inhibitory inputs or the molecular mechanisms underlying these synaptic changes; future studies examining specific synaptic afferents, ion channel regulation, and the respective contributions of tau or amyloid pathology will be critical to clarify mechanism. Finally, this study examined a single early time point in a cross-sectional design; longitudinal investigations tracking the progression of LC dysfunction and its relationship to cognitive decline, and testing whether these functional changes are reversible, will be essential for establishing causality, identifying the molecular mediators of synaptic remodeling, and defining the optimal therapeutic window.

## Conclusion

Overall, our results demonstrate that LC neurons undergo synaptic, intrinsic, and ionic remodeling early in AD pathology, before substantial neurodegeneration, producing a novel decoupling of intrinsic pacemaker activity from input-driven excitability. Rather than a simple hypo- or hyperexcitable state, we propose a nuanced pathological picture in which an early shift toward inhibitory synaptic dominance combines with concurrent loss of voltage-gated K⁺ and Na⁺ conductances, mechanistically explaining both the aberrant, uncoupled spontaneous pacemaking and the restricted capacity for sustained, high-frequency evoked output. Given the LC’s role as the brain’s principal source of noradrenergic modulation, this early breakdown in signal integration offers a mechanistic basis for downstream noradrenergic failure and network instability, and highlights synaptic and ion-channel disruption in the LC as a potential biomarker and therapeutic target for prodromal AD.

## Data Availability

The datasets generated and analyzed during the current study are available from the corresponding author upon reasonable request.

## Acknowledgements

Not applicable.

## Contributions

Baizhen Zhao: Conceptualization, Data curation, Software, Formal analysis, Validation, Investigation, Methodology, Writing -original draft; Morgan Li: Software, Formal analysis, Validation, Writing -original draft; Lingxiao Yi: Validation, Writing -original draft; Jiali li: Conceptualization, Data curation, Supervision, Funding acquisition,

Investigation, Methodology, Project administration, Writing – review and editing.

## Abbreviations

AD: Alzheimer’s disease Aβ, amyloid-β
LC: locus coeruleus
sEPSC: spontaneous excitatory postsynaptic current
mEPSC: miniature EPSC
sIPSC: spontaneous inhibitory postsynaptic current
mIPSCs: miniature IPSC
VGLUT2: vesicular glutamate transporter 2
PSD95: postsynaptic density protein 95
VGAT: vesicular GABA transporter
NE: norepinephrine

